# A weak-interaction model defines the cell-autonomous function of DNA methylation in gastrulation

**DOI:** 10.64898/2026.05.14.725081

**Authors:** Saifeng Cheng, Roni Stok Ranen, Yoav Mayshar, Ofir Raz, Aviezer Lifshitz, Evgheni Casimov, Tal Sokolov, Akhiad Bercovich, Raz Ben-Yair, Oren Ben-Kiki, Netta Reines, Ayelet-Hashahar Orenbuch, Shuailin Du, Jinfang Wang, Ying Zhu, Yonatan Stelzer, Amos Tanay

## Abstract

DNA methylation is essential for mammalian embryonic development, yet its functional implications remain debated. Here, we introduce a quantitative model that reconciles conflicting perspectives on the roles of methylation in genome regulation and cell specification. Using single-cell multiome analysis in chimeric and whole embryo mutants during mouse gastrulation and organogenesis, we separate cell-autonomous from indirect effects of methylation and demethylation machinery. We show that while the transcriptional and chromatin programs defining basic lineages can be established independently of methylation, the fidelity of differentiated states is severely impaired in its absence. Specifically, we identify hundreds of genes and thousands of *cis*-regulatory elements (CREs) dependent on methylation for precise regulation. CRE accessibility alterations in methylation mutants are linked with CpG dinucleotide content, and correlate with multiple transcription factor binding motifs. Our data support a weak-interaction model in which DNA methylation moderates, but does not block or instruct the potency of *trans*-acting regulatory machineries genome-wide.

## Main text

Embryonic development entails the diversification of pluripotent cells through epigenetic and transcriptional alterations. In mammals, this process is accompanied by global remodeling of cytosine methylation, mainly in the context of CpG dinucleotides, rapidly forming a characteristic bimodal methylation landscape: the bulk of the genome comprises low-density CpGs that are methylated by *de novo* methyltransferases (DNMTs) DNMT3A and DNMT3B^1–5^. In contrast, most regions with high-density CpGs, known as CpG islands (CGIs), are protected from *de novo* methylation via the binding of *trans*-factors at these sites^6–8^. Global methylation patterns are copied to daughter cells via a cell-cycle-dependent housekeeping activity of DNMT1^9^. Loss of DNA methylation can therefore occur passively, by incomplete maintenance during cell replication^10^. Alternatively, hydroxymethylation by one of the three ten-eleven translocation (*Tet*) dioxygenase enzymes (TET1/2/3)^11–14^, or binding of *trans*-factors, can induce local demethylation^15^. The importance of DNA methylation to cellular differentiation is emphasized by the observations that mouse embryos deficient in their ability to catalyze^16–20^, maintain^16,17,21–26^, or remove^13,23,27–33^ DNA methylation fail to develop properly.

However, studying how DNA methylation impacts gene regulation during early development remains notoriously challenging for several reasons. First, accumulated data do not support a simple, direct, instructive role for DNA methylation in promoter regulation^34–36^. Instead, methylation is mostly dynamic at distal *cis*-regulatory elements (CREs)^37–40^, where the impact on gene expression is more nuanced and harder to evaluate. Second, the intricate interplay between DNA methylation, chromatin makeup, and *trans*-acting factors is difficult to disentangle^15,41–44^, especially when considering it as a dynamic process with multiple backups rather than a static structure^44^. The correlation between CRE methylation and transcription in embryos is thus hard to interpret functionally, as it is shaped by opposing activities of methylation and demethylation, which may both be affected by parallel epigenetic mechanisms^45^. Lastly, when studying mutant embryos to infer causal effects, the genome-wide impact of alterations in methylation can potentially propagate downstream effects into differentiation catastrophes. This poses great difficulties in analyzing and interpreting the actual cell-autonomous effects of DNA methylation and distinguishing these from indirect effects.

We have recently demonstrated that a single-embryo, single-cell, time-resolved model of mouse gastrulation^46,47^ can serve as a powerful quantitative reference for studying cell-autonomous effects following gene perturbations^46–48^. In this approach, knockout cells develop within the context of a normal embryo that maintains a proper signaling environment, thus reducing the potential non-cell-autonomous effects of the mutation. By analyzing each embryo separately, host and knockout cells from the same embryo can be directly compared, providing a time-matched and controlled system for dissecting gene function. Using this approach, we found *Tet* triple knockout (*Tet*-TKO) cells, which lack the ability for active demethylation, are still capable of differentiating into most gastrulating cell states^48^. This is in contrast to embryos fully composed of *Tet*-TKO cells, which completely failed to develop past this stage^23,27,30^. At the cell-autonomous level, active demethylation mainly affected intermediately methylated regions, which predominantly constitute CREs. Without a functioning TET system, CREs became fully methylated, but this resulted in only small, quantitative effects on gene expression. How these quantitative effects are mediated remains unclear.

Here, we present an enhanced and complete library of knockout models by analyzing mutants with no functional DNA methylation machinery (*Dnmt*-TKO, *Dnmt*-DKO), which are added to the set of TET mutants we introduced before. With this comprehensive collection, we analyze the association between DNA methylation and CRE dynamics by simultaneously measuring transcription and chromatin accessibility profiles in single cells derived from chimeric mutants. These new data provide a long-awaited quantitative affirmation of the classical hypothesis on DNA methylation as a genome-wide secondary regulatory mechanism. It shows that transcription factors (TFs) and their co-factors can establish chromatin accessibility at thousands of ‘strong’ CREs independent of DNA methylation, while accessibility at thousands of additional potential CREs reciprocally correlates with hypo- or hyper-methylation. The data lead to new hypotheses about co-factors interacting with the methylation machinery, suggesting that weak interactions of such factors with CREs and promoters modulate chromatin accessibility and result in pervasive changes in gene expression that are locally subtle but globally impactful.

## Results

### Growth retardation in whole embryo tetraploid *Dnmt* mutants

To systematically dissect the impact of DNA methylation on mouse gastrulation, we added to our previously described collection of TET mutants a double knockout of *Dnmt3A* and *Dnmt3B* genes (*Dnmt-*DKO) and a triple knockout, combined with *Dnmt1* loss-of-function (*Dnmt*-TKO). While both TKO and DKO mutants lose genome-wide methylation, DKO cells retain methylation at parental imprints and retrotransposons^5,17,23,49–51^. We also generated single gene knockouts targeting these genes, with the addition of *Uhrf1* (Extended Data Fig.1a-c), which were used to round up an extensive dataset of single-cell molecular states across the different mutants (Extended Data Fig.1d). We injected GFP-labeled *Dnmt*-TKO and *Dnmt*-DKO cells into tetraploid (4N) blastocysts to obtain mESCs-derived whole-embryo mutants (Fig.1a). Morphological analysis at embryonic day (E) 8.5 showed severe growth delay associated with *Dnmt*-TKO mutant embryos, including lack of observable somites and retarded head fold structure (Fig.1b and Extended Data Fig.2a). *Dnmt*-DKO embryos showed largely normal structures at E7.5. However, the most advanced embryo we recovered recapitulated the previously reported phenotype at E8.5, characterized by a deformed trunk, a relatively large anterior portion, and what appears to be excessive extraembryonic mesoderm in the most posterior portion^16,17^ (Extended Data Fig.2a).

**Fig.1.**
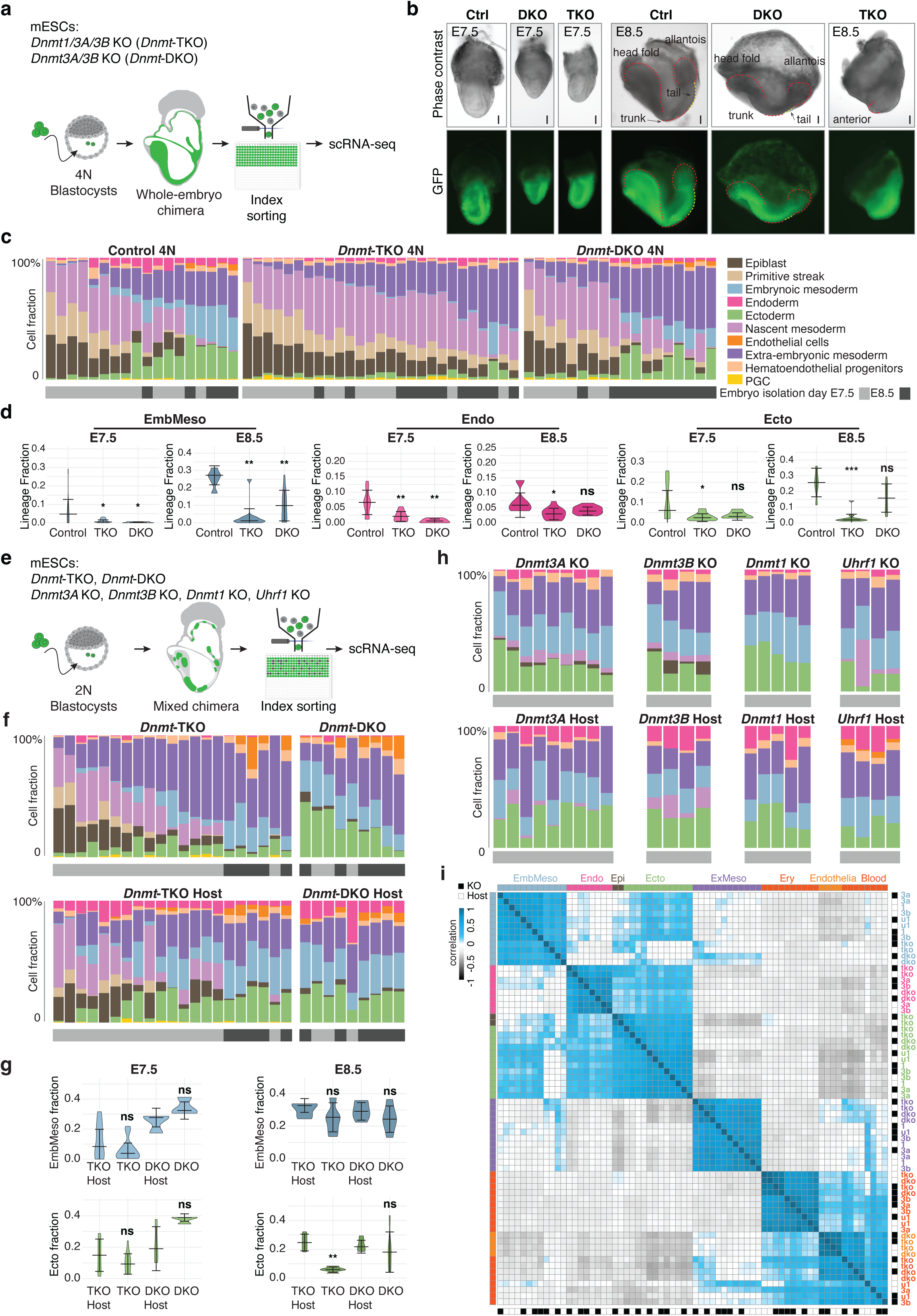
Single-cell quantification of differentiation in DNA methylation mutant embryos. **a**, Schematics showing the generation of whole-embryo mutants by injecting mutant mESCs into 4N blastocysts. **b**, Representative images of E7.5 and E8.5 *Dnmt*-DKO and -TKO whole-embryo mutants. Dashed lines depict embryo structure. Embryos oriented with their posterior end to the right. Scale bars, 100 μm. **c**, Cell-state composition per embryo. Embryos (represented by columns) are arranged according to their proportion of extraembryonic mesoderm relative to their proportion of earlier cell states (epiblast, primitive streak, nascent mesoderm). Isolation day (E7.5, E8.5) for each embryo was labeled in light and dark gray, respectively. **d**, Quantification of the frequency of major lineages per indicated genotype for whole-embryo mutants isolated at E7.5 and E8.5. ns, not significant; *, *p* value < 0.05; **, *p* value < 0.01; ***, *p* value < 0.001. **e**, Experimental design for generating mixed chimeric embryos by injecting mutant cells into normal (2N) host blastocysts. **f**, Cell-state compositions per KO-derived cells (top) or host cells (bottom) are color-coded (columns) as in Fig.1c. Embryo isolation day is depicted as gray/black bars. **g**, Quantification of major lineages for mixed chimeric mutants. Shown are comparisons between the host and the corresponding mutant cell contributions. ns, not significant; **, *p* value < 0.05. **h**, Cell-state composition per embryo as in Figure 1F for single mutant chimeric embryos. **i**, Heatmap matrix showing pooled expression correlation between cell lineages profiled in different genotypes, using either host or knockout cells. Cell lineages are color-coded as in 1C, and host/knockout identification is depicted using black/white – see right and bottom bars. 3a, *Dnmt3A*; 3b, *Dnmt3b*; 1, *Dnmt1*; u1, *Uhrf1*.

### Single embryo analysis shows methylation loss leads to lineage desynchronization

To capture the cellular response to complete DNA methylation loss, we performed single embryo, single-cell RNA-seq on 18 *Dnmt*-DKO and 27 *Dnmt*-TKO mutants isolated from E7.5 to E8.5 (Extended Data Fig.1d, 2b). A total of 9,620 *Dnmt*-DKO cells and 15,567 *Dnmt*-TKO cells passed our quality control. A Metacell (MC) model was used to represent and compare methylation-impaired transcriptional states quantitatively and systematically across embryonic lineages^52,53^ (Extended Data Fig.2c). We analyzed cells from individual mutant embryos by projecting onto a wildtype (WT) temporal gastrulation reference atlas^47^ (see Methods). Mutants exhibited skewed development, with inferred embryonic time of E8.5 embryos poorly reflecting the embryos’ sampling time or expected cell state compositions (Extended Data Fig.2d), in contrast to the highly canonical temporal sequence observed in WT embryos. *Dnmt*-DKO and -TKO mutants exhibit a marked reduction in advanced mesoderm in E8.5 embryos (*p* <0.01, *p* <0.01), and an early and excessive emergence of extraembryonic mesoderm (Fig.1c, d and Extended Data Fig.2e). Consistent with the observed developmental phenotypes, we found a significant decrease in ectoderm cell states in *Dnmt*-TKO embryos (*p* <0.05 for E7.5, *p* <0.001 for E8.5) that was less apparent in *Dnmt*-DKO mutants. We also observed near-complete loss of endoderm cell states in both *Dnmt* mutants (note that host visceral endoderm cells were still observed due to the nature of the tetraploid complementation assay^54,55^). Collectively, morphological and single-cell transcriptional analyses show that disrupting the DNA methylation machinery of the entire embryo leads to growth retardation associated with altered cellular state distributions.

### *Dnmt* mutants regain developmental potency in mixed chimeric embryos

We next used a *Dnmt*-TKO and -DKO chimera assay to test if WT cells, and the signaling they secrete, can rescue some of the developmental abnormalities observed in *Dnmt* whole embryo mutants. When injecting *Dnmt*-DKO or *Dnmt*-TKO mESCs into diploid (2N) blastocysts (Fig.1e), we observed a normal morphology for low to medium levels of chimerism and phenotypes resembling those of whole-embryo mutants in relatively higher levels of chimerism (Extended Data Fig.3a,b). To further understand how mutant cells contribute to this normal-like phenotype, we performed single-cell RNA-seq on individual mixed chimeric embryos isolated from E7.5 and E8.5 time points. For each embryo, we then inferred cell-state compositions separately for KO and host cells. Interestingly, in contrast with whole embryo mutants, *Dnmt*-DKO and -TKO mutant cells within mixed chimeric embryos showed compositional distributions more similar to WT embryos or host cells (Fig.1f and Extended Data Fig.3c). Most notably, we found significantly improved differentiation of TKO and DKO mutant cells toward the embryonic mesoderm lineage (Fig.1g). However, the chimera assays also highlighted potential cell-autonomous effects following methylation loss. Chimeric embryos harboring TKO cells showed a consistent reduction in ectoderm and an increase in extraembryonic mesoderm states compared with the host (Fig.1g and Extended Data Fig.3d). Though very low contribution of *Dnmt* mutant cells to the endoderm was observed from the single-cell RNA-seq data (Fig.1f and Extended Data Fig.3d), immunostaining did show colocalization of mutant with the general endoderm marker FOXA2 in mixed chimeric settings (Extended Data Fig.3e). This effect, may partly be explained by the convergence of extraembryonic endoderm cells into the gut during gastrulation, which introduces a flux of WT cells, but not mutant cells, into this lineage^56–58^. To explore this further, we performed additional experiments with chimeric embryos constructed from knockouts of single methylation machinery genes. Cell composition analysis revealed comparable contributions to all lineages, with a reduced but still evident contribution to the endoderm (Fig.1h and Extended Data Fig.3f).

To unbiasedly assess the potency of the different DNMT mutant lines to differentiate given a normal host signaling environment, we pooled data from knockout cells and their embryo-matched host cells into broad categories of epiblast, ectoderm, endoderm, embryonic and extraembryonic mesoderm, hematoendothelial, primitive erythrocytes, and blood. Systematic comparisons of host- and knockout-derived profiles revealed robust clustering by differentiated state across mutants (Fig.1i). We noted that this conservation of lineage identities is driven by the strongest lineage-specific transcriptional signatures, which may still mask extensive mutant-driven transcriptional perturbations. Therefore, we next aimed at an in-depth quantitative analysis of the fidelity of the entire lineage-specific regulatory programs between mutants and host cells.

### *Dnmt* mutants present quantitatively perturbed gastrulation states

We focused on the earliest gene expression effects of methylation depletion in pluripotent epiblast cells, where aberrant intercellular signaling effects could not yet accumulate to trigger indirect mutant expression changes. To maximize quantitative precision, we excluded genes located on trisomic chromosomes for each mutant line (Extended Data Fig.4a). Comparison of mutant and control epiblast expression was then performed over two axes representing the ‘core*’* transcriptional program and its ‘fine-tuned*’* state. This took advantage of our assay’s single embryo resolution – allowing precise timing of mutant and WT epiblast cells using one signature and controlled analysis of the remaining gene expression variation. We defined the core epiblast program using a transcriptional signature based on genes maximally correlated with the epiblast transcription factor *Utf1* and assessed the fine-tuned state by estimating a transcriptional divergence score between mutants and matched WT embryos (Extended Data Fig.4b; see Methods). This analysis showed that *Dnmt* mutant cells, from either whole-embryo or mixed-chimeric embryos, maintain WT-like core-program expression (Fig.2a). This was in contrast to the fine-tuned mutant epiblast state, which showed high divergence levels compared to controls across multiple single embryos (Fig.2b). In a chimeric context, even WT host cells showed some divergence in comparison with controls, probably due to interactions with mutant cells. Still, this divergence was much lower than recorded for methylation-deprived cells. Similar trends were observed for the more differentiated nascent mesoderm cells, including a conserved core program (in this case, nascent mesoderm-specific genes), along with a high level of expression divergence for genes not within the core program (Fig.2c,d and Extended Data Fig.4b).

**Fig.2.**
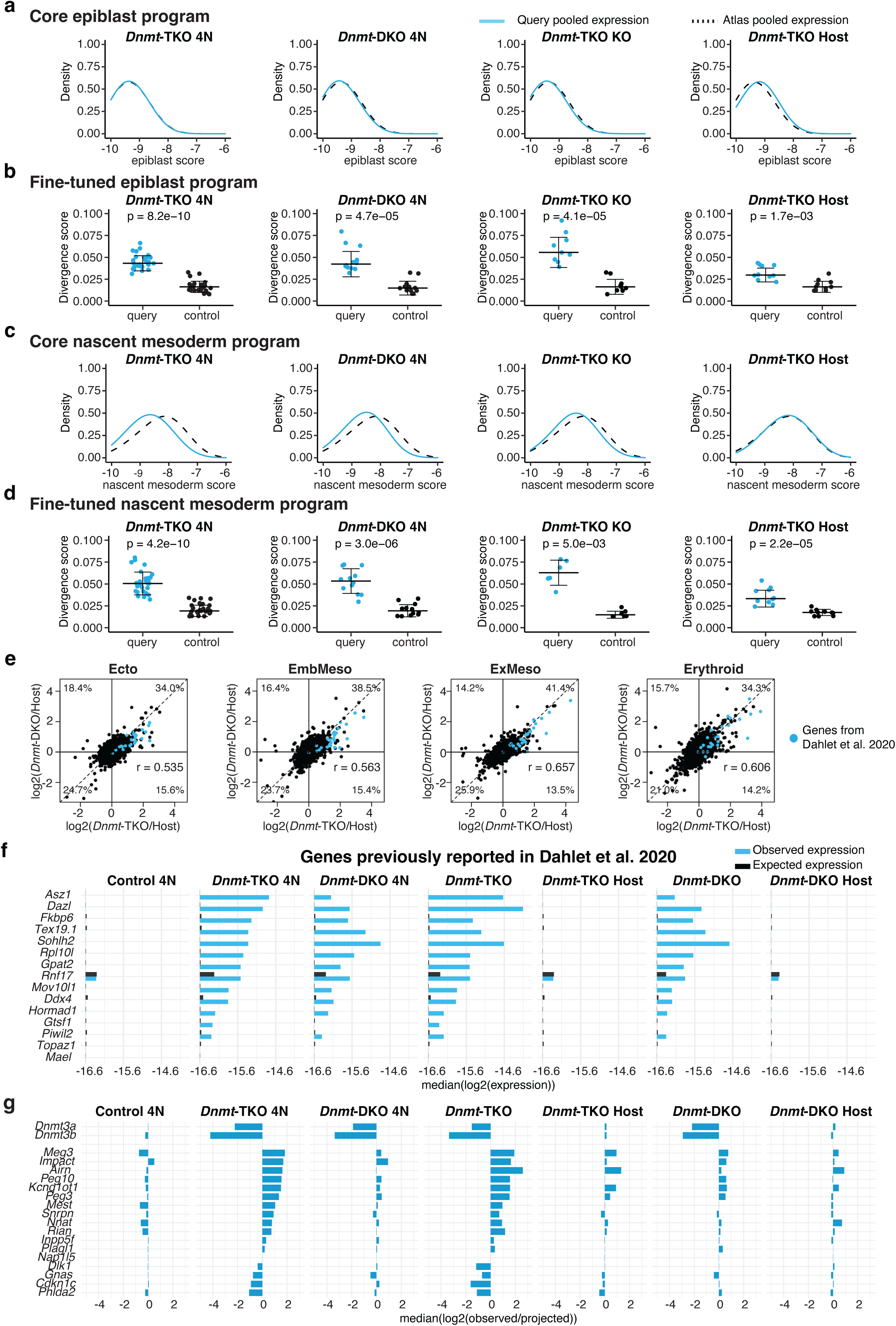
Methylation-dependent transcriptional perturbation over conserved cell states. **a**, Distributions of core epiblast program genes (log2 total normalized expression) among epiblast cells of the indicated genotypes from whole-embryo mutants (4N) and mixed chimeras (see Methods). Distributions of matching atlas embryo references are shown in dashed lines. The X-axis represents absolute expression (log2 of relative unique molecular identifier [UMI] frequency). **b**, Embryo-embryo variance scores (“divergence scores”) for query embryos’ epiblast (KO-derived or host-derived) and their best-matched atlas control embryos’ epiblast; see Methods. **c**, Density plot of aggregated single-cell expression of core nascent mesoderm program genes as described in **a**. **d**, Embryo-embryo variance scores for query embryos’ early nascent mesoderm (KO-derived or host-derived) and their best-matched atlas control embryos’ early nascent mesoderm; see Methods. **e**, Comparison of global gene expression changes between *Dnmt*-DKO and -TKO cells and their respective host cells for the indicated cell lineages. Previously reported methylation-affected genes are highlighted in blue. Percentages in each corner represent the proportion of total genes in each quadrant. r represents the calculated Pearson correlation between genes affected in *Dnmt*-DKO and *Dnmt*-TKO. **f**, Expression of methylation-affected genes from Dahalet et al., 2020 in the indicated genotypes from mixed chimeras and whole-embryo (4N) mutants, compared with WT atlas expression (see Methods). **g**, Expression of imprinted genes in indicated genotypes from mixed chimeras and whole-embryo (4N) mutants, compared with WT atlas expression (see Methods).

### Loss of DNA methylation perturbs transcription consistently between mutants

To test whether removal of DNA methylation results in similar transcriptional de-tuning in DKO and TKO cells, we computed differential expression between mutant and host cells in each profiled embryo and pooled the results across broad differentiation lineages (Fig.2e). Interestingly, this analysis showed conserved methylation-dependent deregulation across lineages (r = 0.535 to 0.657, depending on lineage and its coverage). We could not robustly test the effects in the endoderm, given its very low representation in mutants. Transcriptional perturbation mostly involved gene induction (or derepression), whereas repression was less noticeable. We next screened for genes consistently deregulated in both TKO and DKO cells across different lineages, and identified over 1,000 genes with significant (FDR < 0.01) expression changes in both genotypes and in at least one lineage (Extended Data Fig.5a and Supplementary Table 1). While the extent of derepression varies across lineages, on average 52.6% of affected genes are consistently derepressed. Among these genes were previously reported germline-specific and piRNA targets^17,50^, that are not expressed in WT cells but are induced in *Dnmt*-DKO and -TKO cells (Fig.2f). We noted that, compared with lineage-specific TFs or housekeeping genes, the expression of these methylation-dependent genes is weak in absolute terms, even following derepression in the mutants (Extended Data Fig.5b).

An exceptional set of genes that are expected to be directly affected by DNA methylation are parentally imprinted genes. DNA methylation imprints are set in the germline and are passively maintained following fertilization by the housekeeping activity of *Dnmt1* and *Uhrf1*, ultimately regulating the monoallelic expression of imprinted genes in clusters. Indeed, *Dnmt-*TKO, but not *Dnmt*-DKO cells, showed global alterations in imprinted gene expression, compared with WT atlas or host cells (Fig.2g and Extended Data Fig.5c). This also shows that the consistent transcriptional effects we observed between DKO and TKO are not merely a consequence of loss of imprinting events, as the latter is observed only in TKO. Taken together, single-cell RNA-seq data indicate that transcriptional sorting into the main developmental lineages occurs robustly without DNA methylation. However, the fidelity of gene expression control in methylation-deprived differentiated states is low, with numerous weak but consistent gene expression changes.

### Mutants perturb chromatin accessibility within conserved lineage states

We reasoned that the broad quantitative and cell-autonomous transcriptional perturbation we observed above must be driven, at least in part, by methylation-dependent in-*cis* interactions between gene regulatory machinery and methylated/unmethylated DNA. To search for and characterize such effects, we profiled simultaneously chromosomal accessibility and RNA from single mutant or host cells derived from mixed chimeric embryos (Fig.3a). Data were generated from *Dnmt*-TKO, *Dnmt*-DKO mixed chimeric embryos, as well as *Tet*-TKO embryos, for which we previously demonstrated genome-wide methylation gains^48^. We used the extracted cells’ transcriptional profiles to infer a metacell model that we annotated by comparison to the WT atlas (Extended Data Fig.6a; See Methods). We then pooled ATAC profiles in each metacell and collected metacells sharing similar atlas annotations to create quantitative, normalized cell type accessibility profiles (Extended Data Fig.6b). We separately studied accessibility at transcription start sites (TSS) and putative CREs. The latter were defined *de-novo* as ATAC hotspots not associated with a known TSS. Analysis of the accessibility landscape in WT and mutant cells across broad cell states showed robust clustering of profiles by lineage, rather than methylation state (Fig.3b). This indicated a broad conservation of methylation-independent lineage-specific accessibility profiles. However, when considering profiles normalized over their respective lineage mean, we derived robust clustering of *Dnmt*-TKO and DKO across lineages, with WT profiles and *Tet*-TKO profiles forming separate clusters (Extended Data Fig.6c). Together, this data is consistent with the observation made above for transcriptional states: overall conservation of the basic lineages, with methylation-dependent consistent perturbation within each lineage state.

**Fig.3.**
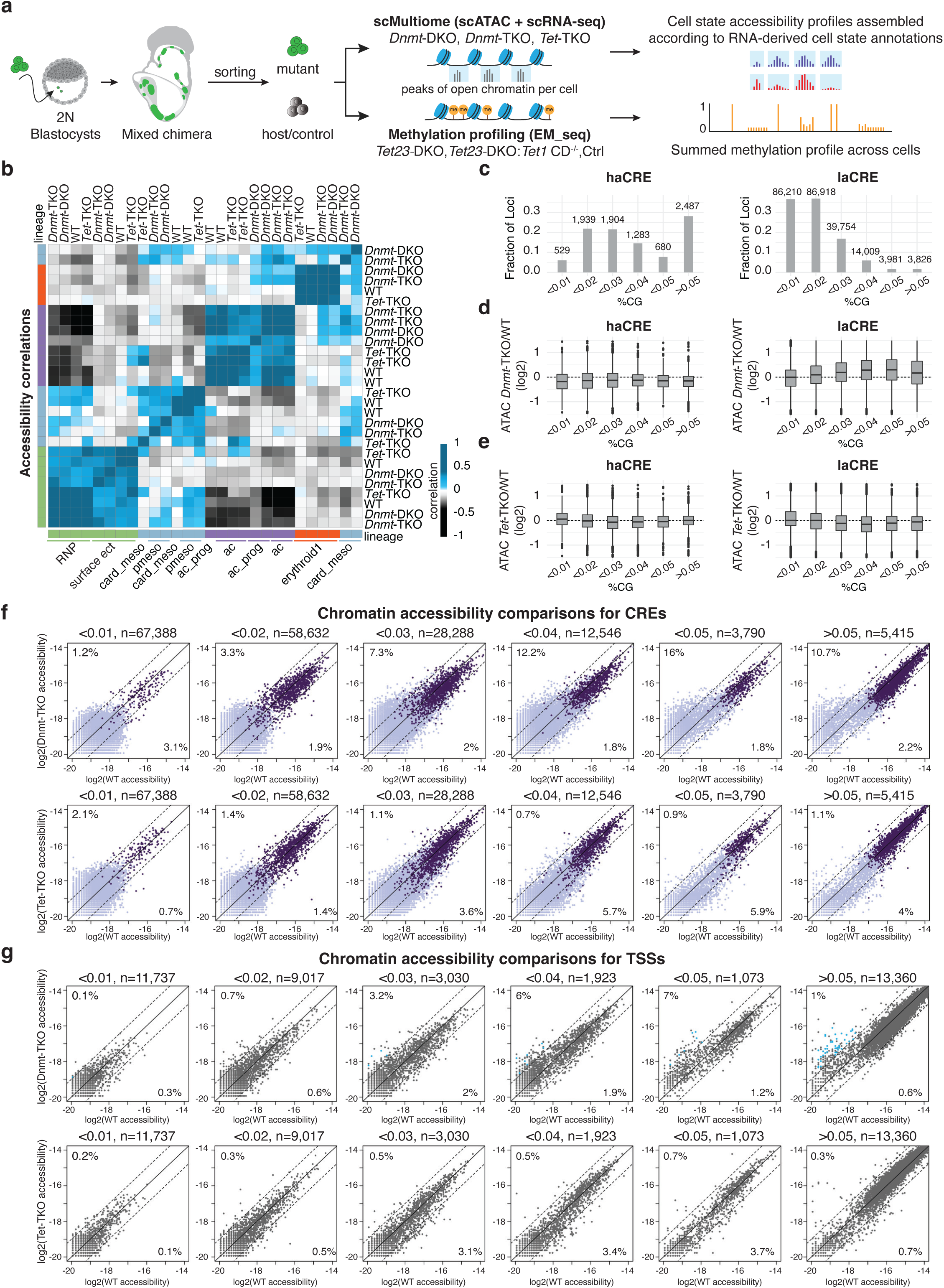
Widespread divergence in weak CRE accessibility in methylation mutant cells. **a**, Epigenomic profiling of chimeric embryos using scMultiome and EM-seq. **b**, Correlation between normalized chromatin accessibility profiles in indicated cell states and genotypes. **c**, Shown are numbers and fractions of CREs that are grouped according to CpG content and normalized accessibility (haCRE: > -16.5, laCRE < -16.5). **d**-**e**, Differential accessibility for CREs across different CpG content bins in *Dnmt*-TKO cells (**d**) and *Tet*-TKO cells (**e**), compared to WT atlas cells. Data are shown for pooled ectoderm profiles. **f**, Depicting *Dnmt*-TKO CRE accessibility vs. WT (top row) and *Tet*-TKO CRE accessibility vs. WT (bottom row). Loci are binned by CpG content. The percentage of loci with either a gain or loss of accessibility per bin (fold change > 1 or < -1) is indicated. haCREs (classified as in **c**) are highlighted in dark purple. **g**, Similar to **f** but showing data for TSSs. Methylation-sensitive genes (Fig.2f) are highlighted in blue.

### Methylation affects accessibility weakly but on a genome-wide scale

Considering that methylation can affect the properties of DNA and chromatin mainly through CpG dinucleotides, we stratified all identified CREs by CpG content and profiled the distribution of accessibility change following methylation loss (in *Dnmt* mutants) or gain (in *Tet* mutants). We further partitioned CREs into high-accessibility elements (*haCRE*s), defined as having WT log relative accessibility exceeding -16.5, corresponding to at least 20% of the maximal accessibility observed at promoters, and low-accessibility elements (*laCREs*), comprising all remaining CREs. haCREs were enriched at high CpG content loci (Fig.3c), and generally showed no methylation or demethylation sensitivity. In marked contrast, laCREs showed pervasive accessibility gain in DNMT mutants, with the effect being particularly pronounced at intermediate CpG content (between 2%-5%) (Fig.3d). The highest median fold change of 0.36 was observed at the 4%-5% CpG content bin (*p* <0.001, compared to the lowest CpG bin). A reciprocal (albeit weaker) effect was observed in *Tet*-TKO mutants (Fig.3e), with the lowest median fold change of -0.14 observed at the 3%-4% CpG content bin (*p* <0.001 compared to the lowest CpG bin). These trends were consistent between ectoderm (Fig.3c-e) and extraembryonic mesoderm cells (Extended Data Fig.6d). Fig.3f shows a more detailed analysis comparing WT and *Dnmt*/*Tet* mutants’ accessibility for all laCREs and haCREs, grouped by CpG content. At the higher end of the accessibility distribution (haCRE elements, dark purple dots), no changes are observed. However, in regions with low WT accessibility (laCRE elements), a remarkable genome-wide accessibility gain is observed, especially in the intermediate CpG content bins. We found that up to 16% of the 4-5% CpG content bin exhibit more than a 2-fold increase in accessibility. But, given their very low basal accessibility, these affected elements remain weakly accessible despite their demethylation-dependent induction. The reciprocal effect in *Tet*-TKO cells was implicated mostly in weakly accessible CREs, with up to 5.9% of the elements in a single bin repressed 2-fold. A similar, albeit weaker, trend was observed when analyzing TSSs with intermediate CpG content (Fig.3g). These results suggest that methylation regulates accessibility at thousands of CREs, typically acting as a weak repressor. At the same time, it also indicates that when strong activation signals are present, methylation has no effect on accessibility.

### Methylation-independent phased nucleosome arrays around CTCF binding sites

So far, we have quantified the pervasive effect of methylation gain and loss on total accessibility at CREs and TSSs. To study the effect at higher resolution, we used a sensitive spatial cluster approach dissecting 2kb accessibility distributions centered on groups of loci of interest, comparing WT, *Dnmt*, and *Tet* mutant profiles. We first applied spatial clustering to the archetypal group of genomic loci targeted by CTCF insulators. These sites are known to anchor a highly organized nucleosome array^59^, which is clearly observed in our normalized accessibility profiles. Importantly, the nucleosome array remains completely unaffected by the loss of either methylation or demethylation (Extended Data Fig.7e,f). We projected WT and *Tet*-TKO methylation on the CTCF spatial clusters and observed that the demethylation in accessibility maxima is unperturbed in TET mutants, but the level of methylation at nucleosome positions (accessibility minima) is increasing (Extended Data Fig.7g, red vs. black). Analysis of spatial clusters at high CpG content (HCG) TSSs showed a similar effect with complete conservation of a broad nucleosome-depleted region (NDR), partial phasing (periodic peaks) of nucleosomes around it, and some shrinkage of the zero methylation regions surrounding these NDRs in TET mutants (Fig.4a-c). Together, the data demonstrated that at CTCF binding sites or high CpG content TSSs, methylation acts downstream of other chromatin-organizing mechanisms. It also showed that demethylation at strong NDRs is TET-independent, whereas methylation at nucleosome positions is affected by TETs.

**Fig.4.**
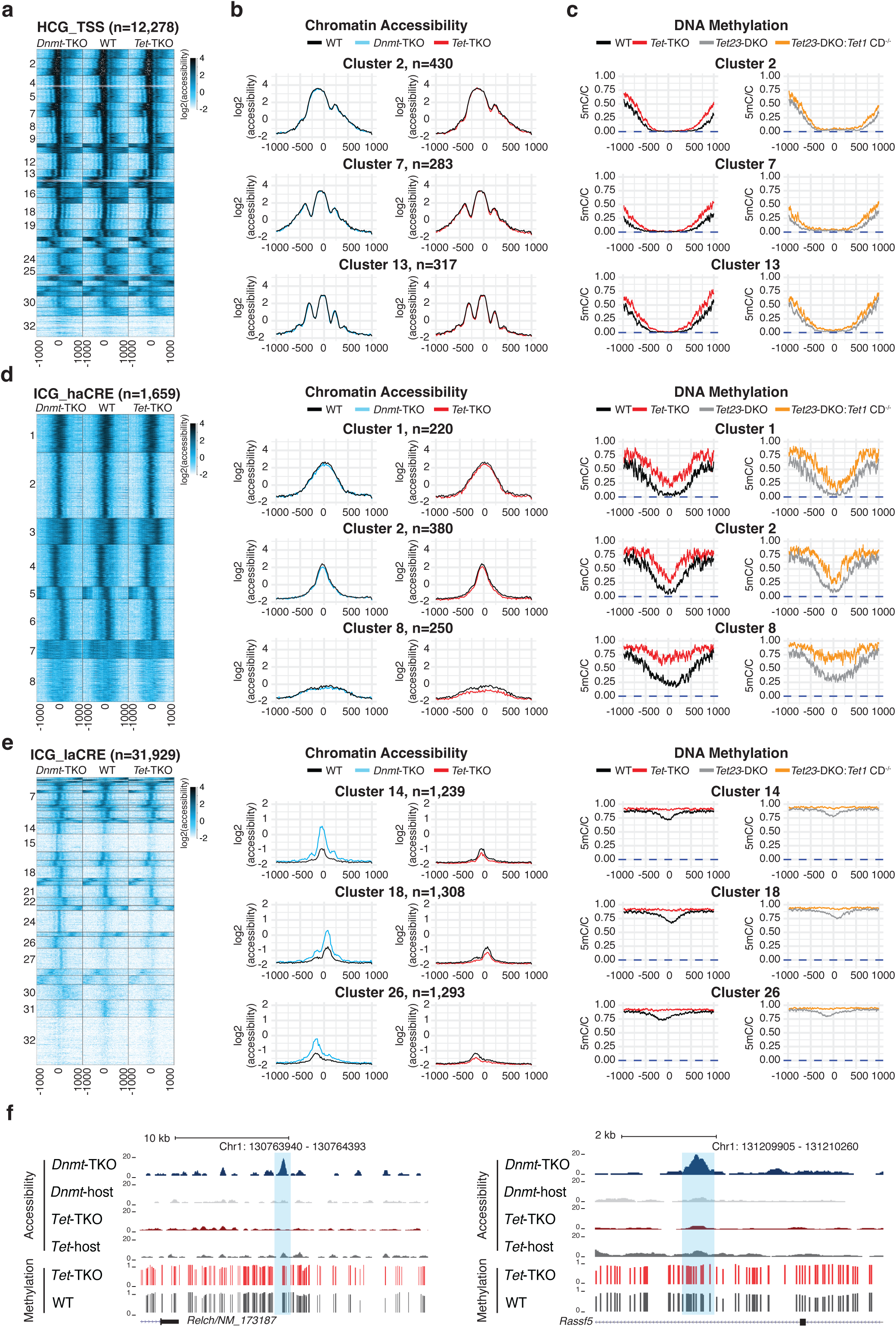
Conservation and methylation-dependent perturbation of chromatin states. **a**, Heatmap of clustered chromatin accessibility profiles for 2kb loci (rows) centered and oriented (left – 5’, right – 3’) according to transcription start sites (TSS) with high CpG density (HCG_TSS, CpG density>0.04). **b**, Quantitative zoom-in for representative clusters from **a**, comparing the three genotypes. **c**, As in **b**, but showing DNA methylation profiles in WT and three different TET mutant lines. The Y-axis represents averaged methylation, ranging from 1 (fully methylated) to 0 (unmethylated). Blue dashed lines indicate the zero expected methylation levels for *Dnmt*-TKO. **d**, **e**, Chromatin accessibility and DNA methylation profiles for high accessibility (**d**) and low accessibility (**e**, all others) CREs with intermediate CpG content (ICG, 0.02 < CpG density < 0.04). **f**, Genome browser tracks of chromatin accessibility for representative ICG_laCREs loci. DNA methylation profiles are calculated for pooled E7.5 WT and *Tet*-TKO embryonic cells.

### The TET1 catalytic activity drives hypomethylation around accessible hotspots

The observed TET-dependent demethylation at nucleosome positions around CTCF could be mediated via TET catalytic activity or by direct DNA binding, which may prevent *de novo* methylation by the DNMTs. To discern between these two alternatives, we have generated mESCs depleted for *Tet2* and *Tet3*, but harboring intact *Tet1* (denoted *Tet23*-DKO), as well as an isogenic cell line in which we mutated the catalytic domain of *Tet1* (termed *Tet23-*DKO; *Tet1-*CD) (Extended Data Fig.7a-d; See Methods). Both genotypes were injected into 2N blastocysts, followed by bulk whole-genome DNA methylation profiling of mutant cells derived from mixed-chimeric embryos at the late headfold stage. We identified remarkably similar methylation patterns in *Tet1*-CD in comparison to *Tet*-TKO mutants, as well as comparable patterns between *Tet23*-DKO and WT control across all genomic contexts examined (Fig. 4c-e and Extended Data Fig.7g,h), ruling out non-catalytic effects of TET DNA binding. Together, we conclude that in CTCF-bound nucleosome arrays and HCG TSSs, it is the catalytic activity of TET1 that attenuates methylation over nucleosomes, while TET-independent mechanisms reduce methylation between the nucleosomes.

### Accessibility is perturbed at ICG loci with high WT methylation

We hypothesized that the genome-wide anti-correlation between accessibility and methylation gain or loss at CREs with intermediate CpG (ICG) content is driven by a balanced interplay between methylation and *trans*-acting factors that repress or promote accessibility. Spatial clustering of ICG CREs with high WT accessibility (n=1,659) revealed robustly conserved accessibility profiles in mutants, showing that at these loci, *trans*-acting factors dominate over DNA methylation (Fig.4d). We found WT methylation at these loci approaches zero at the NDR center, while protection is reduced in *Tet*-TKO cells that develop partial (but not high) methylation. This gain of methylation is shown to be dependent on the TET catalytic activity. In contrast, loci with low WT accessibility (n=31,929, Fig.4e,f) exhibit a localized increase in accessibility in *Dnmt*-TKO and a slight decrease in *Tet*-TKO, along with high DNA methylation levels in WT that reach complete levels in *Tet*-TKO. According to these data (see also data on low CpG content loci, Extended Data Fig.7h), methylation appears inconsequential, functioning downstream of the most potent activating machineries. However, it exerts a pervasive effect at loci engaged with seemingly weaker *trans*-factor activities, characterized by near-complete methylation in WT cells. Importantly, these effects are generally dependent on the availability of CpG sites at the loci. In light of this insight, one must define the interplay between DNA methylation and *trans*-acting factors over a spectrum of CpG densities.

### Multiple TF binding motifs are correlated with methylation-sensitive accessibility

We used a library of candidate TF binding motifs to systematically screen for CRE sequences correlated with accessibility change in *Dnmt* or *Tet* mutant (See Methods). To isolate the motif-dependent effects while controlling for the strong overall trend linking CpG content with methylation-dependent accessibility (i.e., Fig.3c), CREs were analyzed in numbered bins representing defined CpG density ranges (0-1%,1-2%,…,4-5%, >5%). This analysis was performed separately in three cell states, under the assumption that distinct activity of *trans*-factors within these states may influence the links between motifs and accessibility. For each motif, we identified the top (2%) matching CRE sequences and screened for models showing significant enrichments (*q* <0.0001; See Methods) of accessibility gain in *Dnmt*-TKO and/or loss of accessibility in *Tet*-TKO. We grouped motifs (some of which are variants of similar binding preferences) into clusters (Extended Data Fig.8a), and analyzed trends of *Dnmt*-TKO*/Tet*-TKO accessibility changes (Fig.5a; blue and red color coding, respectively, and Supplementary Table 2), basal WT accessibility (green color coding), and changes in methylation between WT and *Tet*-TKO (orange color coding).

**Fig.5.**
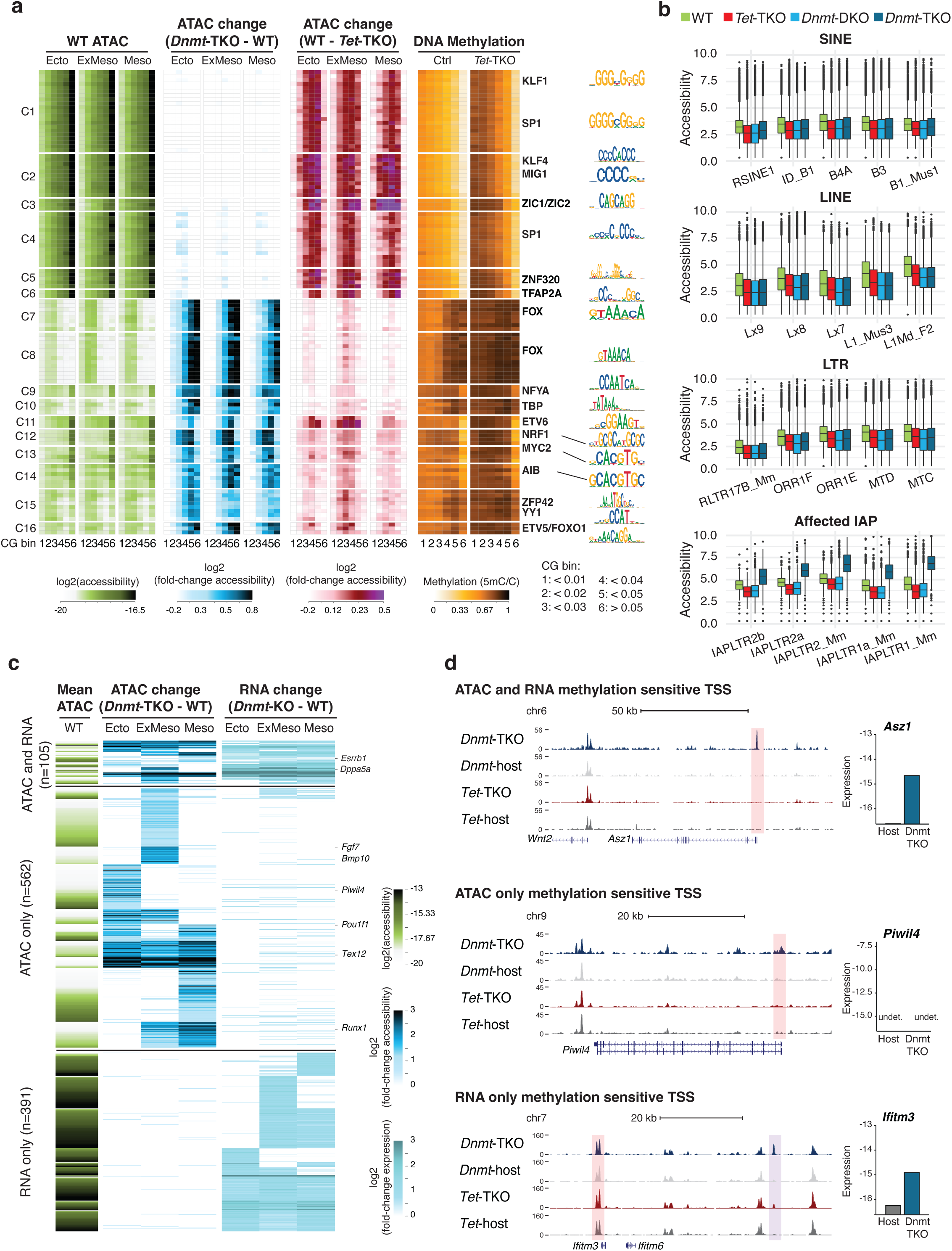
Linking TF binding motifs with methylation-dependent regulation. **a**, We identified TF motif models with statistically significant linkage with methylation-dependent accessibility. Each row represents a subset of CRE loci with high binding affinity for one such TF model (see Methods). Colored-coded matrix depicts WT accessibility, *Dnmt*-TKO accessibility gain, *Tet*-TKO accessibility loss, and DNA methylation across CpG content bins 1 (0-1%) to 6 (>5%). Data are shown for the ectoderm (Ecto), mesoderm (Meso), and extraembryonic mesoderm (ExMeso) lineages. Representative TF model names and their motif logos are shown to the right. **b**, Shown is the logged normalized total chromatin accessibility of transposable elements subfamilies in the indicated genotypes. Total accessibility is represented by the sum of reads from all sequenced cells. **c**, Showing clusters of TSSs with methylation-dependent accessibility and/or expression regulation. Mean WT TSS accessibility is shown for reference (left columns), and the differential accessibility and transcription in *Dnmt*-TKO compared to WT cells (pooling over three lineages) is used to group and cluster TSSs. Complete data is available in Supplementary Table 3. **d**, Genome browser tracks of chromatin accessibility in ectoderm cells of indicated genotypes for representative TSSs from **c**. Absolute expression in host and *Dnmt*-TKO cells for the indicated genes is shown in barplots.

Overall, the impact of methylation on motif-harboring CREs varied according to their WT basal accessibility level. At the lower end of this spectrum, clusters 7-10 showed markedly low WT accessibility levels and high WT methylation. Motifs in these clusters are high AT-content (e.g., of the FOX and NFY family) and are associated with accessibility gain in *Dnmt*-TKO, which is consistent among ectoderm and mesoderm lineages. Clusters 11-16 showed intermediate WT accessibility, and could therefore support both *Dnmt*-TKO-dependent accessibility gain and *Tet*-TKO-dependent accessibility loss. Motifs in these clusters represent exciting potential mechanisms for tunable accessibility switches. For example, cluster 11 (ETS-like motifs) showed preferential methylation-dependent repression in the extraembryonic mesoderm, while cluster 15 (YY1/2 motifs, ZFP42) showed bias toward ectoderm. Other motifs showed a more constitutive effect, for example the previously implicated NRF1 and MYC motifs^60,61^. Interestingly, we note that NRF1 was also observed at low-to-intermediate CpG content TSSs (Extended Data Fig.8b). Finally, motifs in clusters 1-6 were highly accessible in WT, but showed some *Tet-TKO*-dependent accessibility reduction. The models in these clusters are variants of high GC content motifs (e.g., Sp1, Klf, Zic2), with a methylation effect that is consistent among lineages.

It must be noted that the impact of both gaining and losing DNA methylation is limited in intensity. The most active CREs are unlikely to be repressed completely or activated *de novo* when methylation is perturbed. Moreover, only some of the motifs correlated with methylation-dependent accessibility involved a CpG site intrinsically, suggesting that much of the TF-specific effect observed may be driven by complex interactions between co-factors, TFs, and DNA, rather than direct regulation of TF binding. Nevertheless, the data strongly show that methylation affects dozens of factors and thousands of CREs, and that it is likely contributing to mechanisms restricting the non-specific activation of functional CREs in inappropriate lineages.

### Specific accessibility derepression of IAP elements in *Dnmt*-TKO cells

DNA methylation has been shown to play a role in the epigenetic silencing of transposable elements (TEs), and it has been suggested that this function may underlie the evolution of the DNMT machinery and its knockout phenotype^62,63^. We profiled the total accessibility of TE families using the raw multiome reads (see Methods), comparing WT, *Dnmt*-TKO, *Dnmt*-DKO, and *Tet*-TKO pooled cells across all lineages (Fig.5b). No effect was observed for families of LINEs, SINEs, or LTRs, which consist almost exclusively of inactive TEs in the mouse. Remarkably, accessibility analysis of IAP families revealed a significant increase in *Dnmt*-TKO cells, but not in *Dnmt*-DKO or the other lines. This result is consistent with previous studies showing that DNMT1 is sufficient for the maintenance of TE silencing^26^. The robust maintenance of IAP silencing also shows that the consistent TKO/DKO methylation knockout effects we observed above are not driven by derepression of IAPs.

### Association between deregulated expression and accessibility in *Dnmt* mutants

The transcriptional and accessibility methylation effects we described above were analogous, both demonstrating the conservation of basic lineage programs and extensive perturbation of the regulation within them. To study the association between the two effects, we clustered TSSs given differential expression and accessibility in pooled cells from three lineages. This allowed the definition of a group of 105 TSSs in which accessibility gain and transcriptional derepression were jointly observed (Fig.5c,d and Supplementary Table 3). Misregulation of these promoters was consistent between *Dnmt*-TKO and *Dnmt*-DKO, suggesting a direct linkage with methylation changes (Extended Data Fig.8c). Interestingly, among the implicated derepressed genes, we identified the pluripotency TFs *Dppa5a* and *Esrrb* (the latter showing transcriptional derepression only in ectoderm). In a larger number of TSSs, accessibility gain was not accompanied by transcriptional change (middle panel; n=562, Supplementary Table 3), occurring consistently across lineages in most cases and, in some instances, within a specific lineage. These accessibility changes, which involve key developmental loci such as *Pou1f1*, *Tex12*, *Fgf7*, *Bmp10,* and *Runx1*, may not lead to transcriptional changes since the required *trans*-acting machinery is not yet in place. Alternatively, they might be compensated for by alternative TSSs for the same genes (58.2% of genes in this group have alternative TSSs whose accessibility remains undisturbed). In contrast, a third group of TSS (lower panel; n=391, Supplementary Table 3) exhibited transcriptional induction without corresponding changes in accessibility. These loci were, however, already highly accessible in WT cells (Fig.5c,d, green scale color coding). This suggests that their transcriptional regulation may be driven indirectly by interacting CREs, or reflect more nuanced links between methylation, transcription, and accessibility. Yet our analysis of the linkage between CRE accessibility changes and TSS derepression revealed no spatial association to support methylation effects mediated by enhancer-promoter contacts (Extended Data Fig.8d). Nonetheless, such effects are likely responsible for at least some of the transcriptional perturbations observed following methylation loss.

### Propagation of early methylation-dependent effects into organogenesis

By E9.5, with the onset of organogenesis, *de novo* DNA methylation activity is reduced. This provides context to test whether the quantitative, cell-autonomous effects observed during gastrulation persist or become accentuated in later differentiation, where methylation dynamics are stabilized, reaching genome-wide peak levels. Using a newly derived *Dnmt*-DKO clone (Extended Data Fig.9a,h), we repeated the chimeric embryo assays and observed a correlation between the extent of chimeric contribution and developmental outcome. Higher contributions of *Dnmt*-DKO cells were associated with increasingly severe phenotypes, characterized by a pronounced developmental arrest affecting all parts of the embryo (Extended Data Fig.9b). To maximize coverage and resolution, we dissected embryonic head parts from 2 chimeric embryos with a knockout contribution of 37.2% and 52.1%. Cells pooled from these embryos were sorted into host and mutant fractions, followed by multiome analysis, deriving 5,493 and 5,655 single cell profiles, respectively (Fig.6a). Remarkably, transcriptional analysis showed that even at E9.5, methylation loss still permits differentiation into the major embryonic lineages, including ectodermal head cell types and a subset of head mesoderm states (Fig.6b-c). However, cell state composition in the head became skewed between WT and DKO, with neural crest cells consisting almost entirely of WT cells, and mesodermal somitic and sclerotome cells consisting mainly of DKO cells. Differential expression analysis showed that most genes consistently maintain normal expression levels within their respective cell state, even when the representation of these states is skewed (Extended Data Fig.9c). Transcriptional perturbation observed on top of this basal conservation includes many lineage-specific effects (Fig.6d, Extended Data Fig.9d) in addition to a lingering, weak overexpression of the methylation-sensitive “germline” signatures (Extended Data Fig.9e). Together, these observations are consistent with earlier gastrulation-stage analyses, indicating that subtle fine-tuned misregulation can bias lineage representation, without disrupting core transcriptional states or lineage identities.

**Fig.6.**
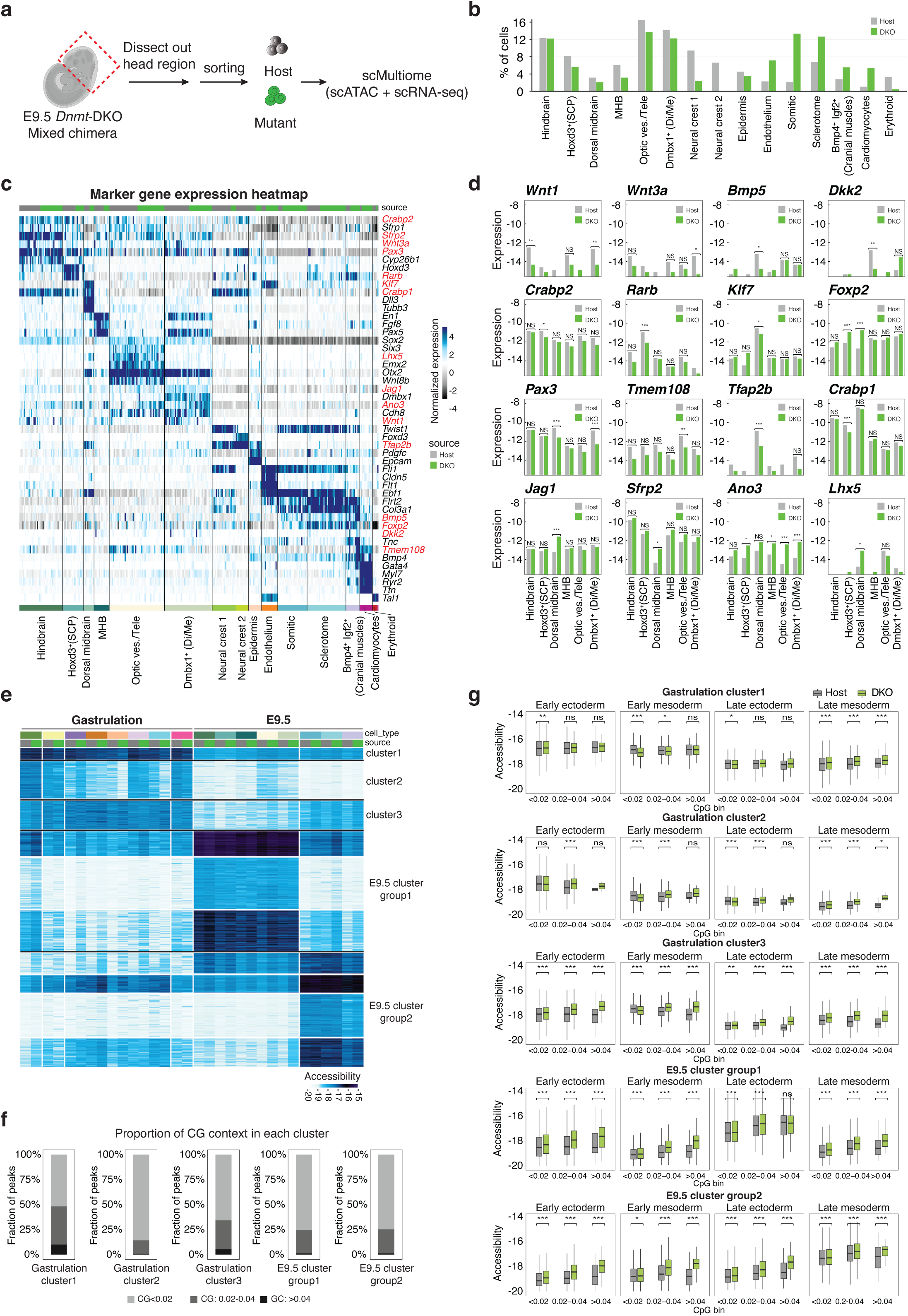
Early embryogenesis embryos exhibit propagated defects of methylation deficiency. **a**, Experimental scheme for E9.5 scMultiome assay. **b**, Relative contributions of each genotype to cell state combination to the total pool of sequenced cells. Hoxd3^+^ (SCP): Hoxd3^+^ spinal cord progenitors; MHB: midbrain-hindbrain boundary; Optic ves./Tele: optic vesicle/telencephalon; Dmbx1^+^ (Di/Me): Dmbx1^+^ diencephalon/mesencephalon. **c**, Heatmap showing marker gene enrichment (expression relative to the median across metacells) across metacells (columns), with genes shown in rows. Metacells are colored by annotated cell states (bottom) and Host/*Dnmt*-DKO genotype (top). Genes whose DGE is highlighted in **d** are marked in red. **d**, Absolute gene expression levels for select genes in Host/*Dnmt*-DKO genotype across six ectoderm cell states. **e**, Clustered heatmap showing chromatin accessibility profiles across pooled cell states per lineage (see annotations in **c**). Loci were selected based on their preference for a specific cell state in host cells. Clusters are grouped (right) based on lineage and developmental-stage preferences. **f**, For each cluster group defined in **e**, CpG content bin composition is displayed as a stacked bar plot representing percentages of peaks belonging to the group. **g**, For each cluster group defined in **e**, accessibility distributions for mean host and mean *Dnmt*-DKO profiles is examined across four panels representing: early ectoderm (rostral neural plate), early mesoderm (ExE mesoderm, allantois, hematoendothelial progenitors, late nascent mesoderm, cardiopharyngeal mesoderm), late ectoderm (hindbrain, Hoxd3^+^(SCP), MHB, Optic ves./ Tele, Dmbx1^+^ (Di/Me)), and late mesoderm (somatic, sclerotome, Bmp4^+^ Igf2^+^ cranial muscles). Accessibility distributions are stratified by CpG content bin. Statistical significance (paired Wilcoxon test between host and *Dnmt*-DKO accessibility) is indicated above each comparison using asterisks.

Analysis of the accessibility landscape across the integrated E9.5 and gastrulation datasets showed clear lineage-specific conservation between the WT and DKO states. This allowed clustering of CREs into groups with similar developmental accessibility trends (Fig.6e, additional clusters shown in Extended Data Fig.9f). Comparative analysis of WT and *Dnmt*-DKO accessibility trends within clusters revealed that loci accessible during gastrulation were at least partly repressed upon later lineage commitment, even in the absence of DNA methylation (Gatrulation clusters 1-3). Reciprocally, aberrant early activation of loci specific to organogenesis was observed in gastrulation states (E9.5 cluster groups 1,2). In general, de-repression tends to occur in contexts where WT accessibility is low (Extended Data Fig.9g). While the majority of cluster CREs were of low CpG content (<2%, Fig.6f), de-repression was confined to CREs with intermediate (>2%) CpG content, consistent with a protective role for DNA methylation at CREs (Fig.6g). Collectively, these data support the weak interaction model, in which lineage transcription factors are required for establishing cis-regulatory accessibility, while DNA methylation modulates their engagement at weak CREs rather than blocking their functional output altogether.

## Discussion

The function of DNA methylation in post-implantation mammalian development is subject to a long-standing debate^35^. On the one hand, the global methylation remodeling at this stage, the severe developmental phenotypes of DNMT and TET mutants in mice, and the well-established role of methylation in maintaining parental imprinting and transposable element silencing, all support its functional importance in securing robust gene regulation. On the other hand, the original hypotheses on the direct and instructive role of DNA methylation in gene repression were largely refuted in functional studies, suggesting that some or even most of the links between DNA methylation dynamics and gene regulation are correlative rather than causal^64–68^. The data we present here reconcile the two seemingly opposing views by putting forward a weak-interaction model for DNA methylation and gene regulation.

We leverage the sensitivity of an experimental system combining single-cell multiomic analysis with chimeric mouse embryos, where either the methylation or demethylation machineries are perturbed (summarized in **Fig 7**). In accordance with models positing low regulatory function of methylation, we first find that mutant cell differentiation into most lineages is feasible when WT host context is co-developing. Nevertheless, aligning with a driving regulatory role for methylation, we identify hundreds of genes and thousands of *cis*-regulatory elements affected by methylation. Accessibility perturbation was examined across different genomic classes, stratified by CpG content and WT accessibility, enabling the detection of subtle effects that would otherwise be difficult to identify. Yet, even when transcriptional or accessibility perturbations are observed, they appear to introduce weak effects rather than complete gene activation or repression. The data show methylation is incapable of protection from *de novo* acquisition at a potent enhancer, in a cell state where active *trans*-acting machinery is available. An in-depth analysis of the associations between TF binding motifs, CpG content, and methylation-dependent accessibility provides robust support for this perspective. A substantial subset of binding motifs is potentially methylation-sensitive, but this sensitivity occurs at loci that are initially inaccessible and develop notable, albeit low, accessibility following loss of methylation.

**Fig.7.**
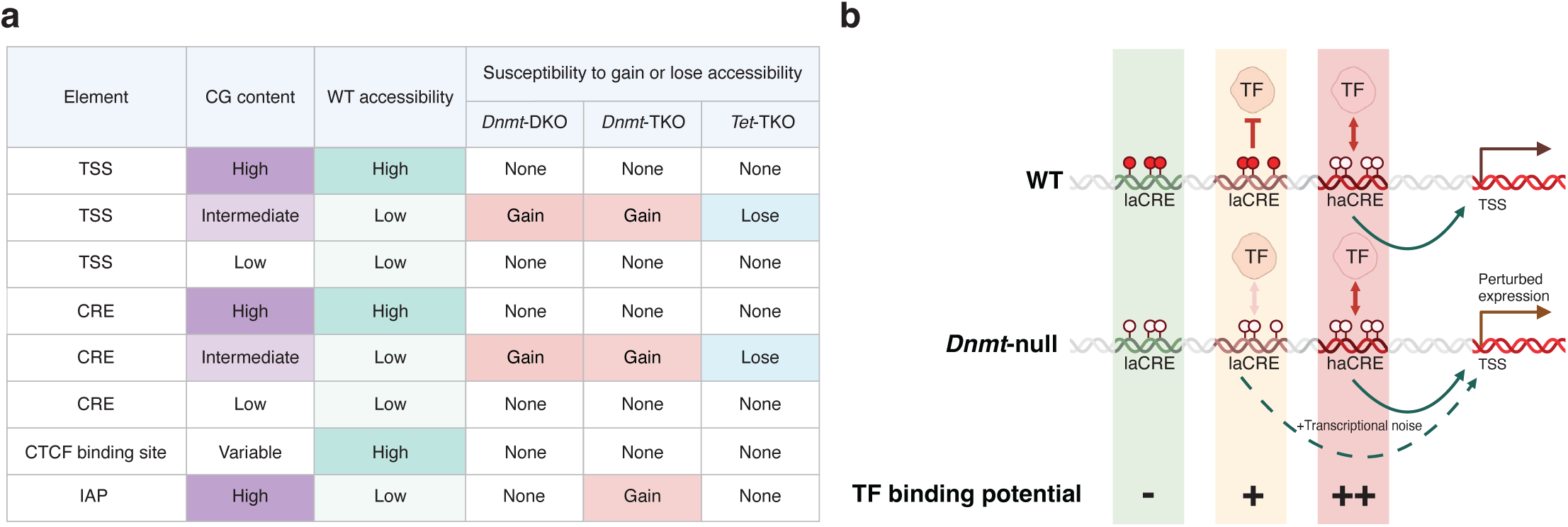
Summary of findings and a schematic illustrating the proposed weak-interaction model. **a**, Table summarizing general observations across genomic categories, highlighting those that are broadly susceptible to changes in chromatin accessibility in the absence (*Dnmt*-DKO and *Dnmt*-TKO) or excess (*Tet*-TKO) of DNA methylation. **b**, Schematic of proposed model. Light orange and pink color highlight columns represent CREs with potential TF binding in the cell, whereas the green highlight column represents a CRE lacking this potential due to the absence of a relevant TF with sufficiently high binding affinity.

A weak-interaction model can provide an attractive basis for understanding how a genome-wide and low-specificity epigenetic mechanism can become critical for normal development and cellular homeostasis. As cells differentiate, their fates become dependent on the robust and specific activation of lineage-determining CREs and downstream genes, primarily through targeting by precise combinations of *trans*-acting factors. Yet there are hundreds of thousands of other elements possessing the latent potential to be activated weakly and spuriously by specific sequence signals (i.e. not a mere stochastic activation of any CRE). Methylation is suggested by our data to provide a (thin) layer of protection against such spurious effects (Fig.7b), acting across thousands of elements through a non-specific and non-instructive mechanism, which is essential for establishing precise cellular states. This is reflected by the fact that, while broad lineage gene regulatory programs are established in the absence of methylation, their fidelity is low and marked by numerous perturbations compared to the WT state.

The well-established functions of DNA methylation in regulation are validated here. Most notably, direct regulation of both IAP and imprint maintenance by *Dnmt1* is documented in *Dnmt*-TKO cells. But as *Dnmt*-DKO cells retain the ability to maintain methylation and silencing at these loci, yet show similar other regulatory effects to *Dnmt*-TKO cells, our data show that the pervasive weak-interaction model is in effect independently of IAPs and imprinting. The behavior of *Tet*-TKO mutants is generally reciprocal to that observed in *Dnmt* mutants. However, at loci protected from WT methylation, no *Dnmt* effect is observed, while at WT loci with near-complete methylation, the impact of TET-mediated demethylation loss can only be minor.

Much remains to be explored in order to understand how the mechanisms characterized here generalize from the very early cell lineages following gastrulation to further refined cell states during organogenesis and in terminally differentiated cells. It is possible that the highly dynamic epigenetic landscape defining the transition from pluripotency to gastrulation modulates the function of DNA methylation toward weaker interactions. More mature cell states may then either increase or decrease their dependency on a properly established methylation landscape, reflecting the activities of DNMT and TET machineries. With the new tools and quantitative models established here, together with the further integration of additional epigenetic layers of regulation, we believe that a fully quantitative model placing DNA methylation in its correct functional and regulatory context is within reach.

## Acknowledgements

We thank the Tanay and Stelzer group members for discussion and advice. Y.S. is the incumbent of the Louis and Ida Rich Career Development Chair and is supported by European Research Council (ERC_CoG EmbryoCellEnsemble), ISF (2824/24), the Minerva Foundation, Helen and Martin Kimmel Stem Cell Institute, and research grant from the Estate of Hermine Miller. S.C. was supported by the EMBO long-term fellowship (ALTF 268-2018). Research in A.T. lab is supported by the European Research Council (ERC cells2Tissues), the Israel Science Foundation BRG program. Research in S.C. lab was founded by the National Natural Science Foundation of China (Grant No. 32570691).This research was further supported Israeli Council for Higher Education (CHE) Data Science program and by a grant from Madame Olga Klein-Astracha.

## Author contributions

S.C., R.S., Y.S. and A.T. conceived the experiments and performed data analysis and interpretation. S.C. generated KO cell lines and collected single-cells with the assistance of R.S., Y.M., and R.B. N.R. prepared scRNA-seq libraries. O.R., A.L., E.C. T.S, A.B., and O.B. assisted with scRNA-seq, multiome, and EM-seq analyses. A.-H.O. and Y.M. performed chimera injections. S.D., J.W., Y,Z., helped in E9.5 multiome assay. S.C., R.R., Y.S. and A.T. wrote the manuscript with input from all the authors.

## Extended Data Figure legends

**Extended Data Fig.1.**
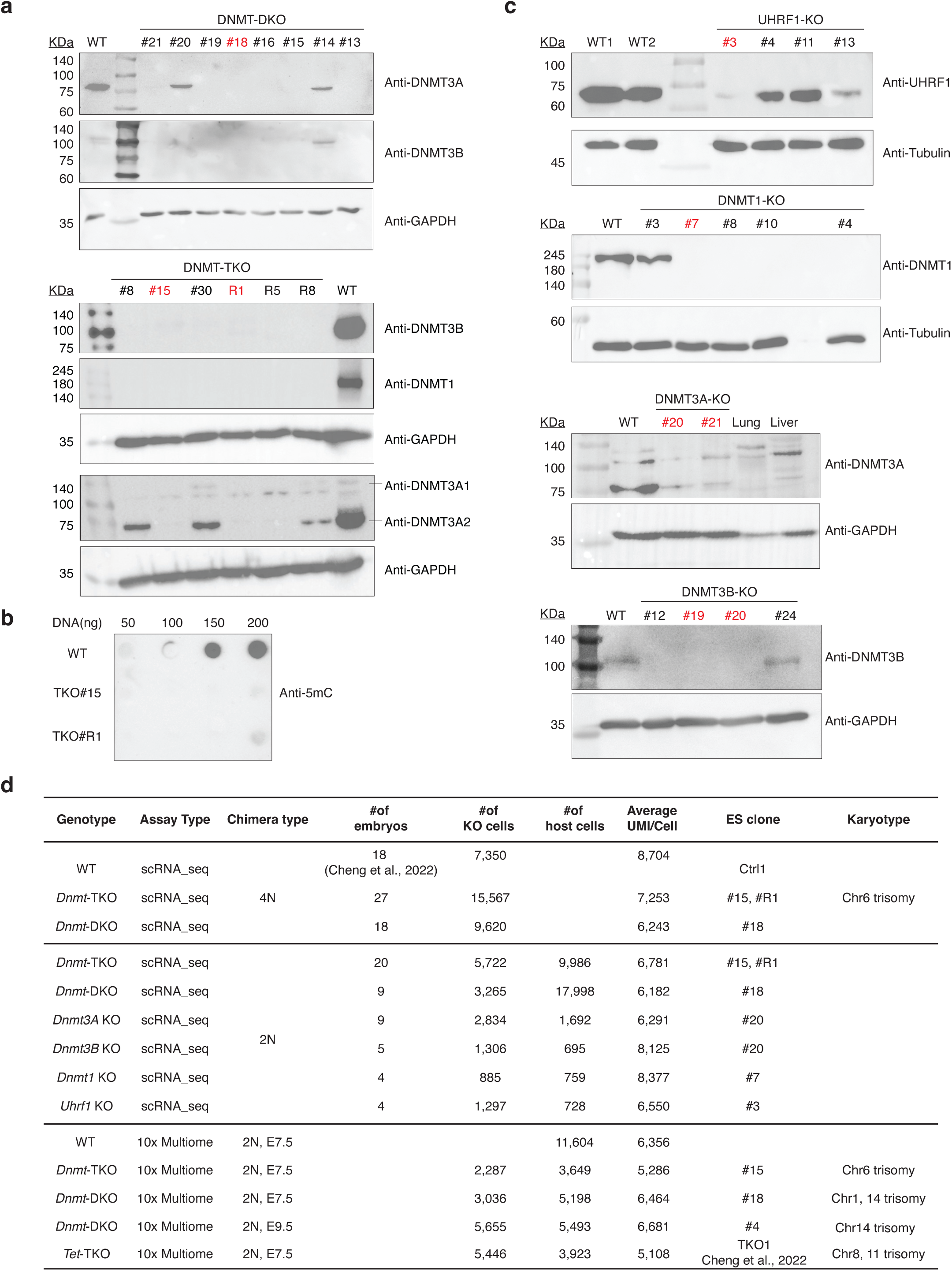
Generation of fluorescently labeled KO mESCs clones for DNA methylation machinery. **a**, Western blot validation for *Dnmt*-DKO and -TKO mESCs clones. Clones labeled in red were used for subsequent chimeric injections. **b**, Dot blot analysis with antibody against 5mC performed on *Dnmt*-TKO clones. **c**, Western blot validation for DNMT1, DNMTA, DNMT3B, and UHRF1 KO mESCs clones. Clones labeled in red were used for subsequent chimeric injections. **d**, Overview of datasets included in this study.

**Extended Data Fig.2.**
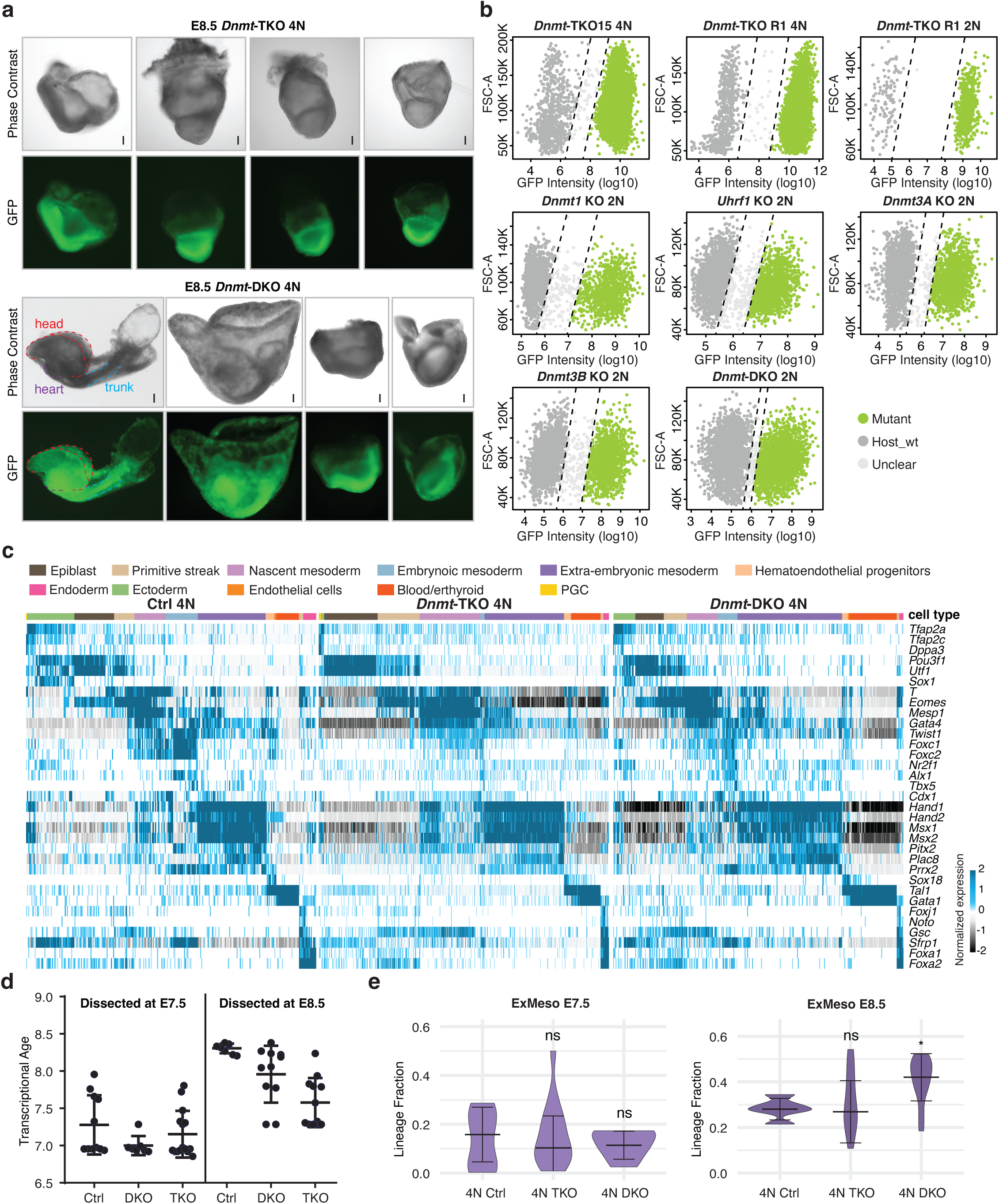
*Dnmt*-TKO and -DKO differentiation in the context of whole-embryo mutants. **a**, Representative images of *Dnmt*-TKO and -DKO whole-embryo mutants at E8.5. Dashed lines depict embryo structure. All embryos are oriented with the anterior part to the left. Scale bars: 100 μm. **b**, Representative FACS plots illustrating the gating strategy for chimeric embryos based on recorded GFP intensity. **c**, Heatmap showing enrichment (expression relative to the median across cell states) of marker gene expression (rows) across metacells (columns) from indicated genotypes, colored by annotated cell states (top). **d**, Deviation of the inferred transcriptional age of each *Dnmt*-DKO and -TKO whole-embryo mutant from their embryonic day (E) dissection time. **e**, Quantification of extraembryonic mesoderm (ExMeso) frequency for Control, *Dnmt*-TKO, and *Dnmt*-DKO derived whole-embryo mutants (4N) isolated at E7.5 and E8.5. ns, not significant; *, *p* value < 0.05; **, *p* value < 0.01; ***, *p* value < 0.001.

**Extended Data Fig.3.**
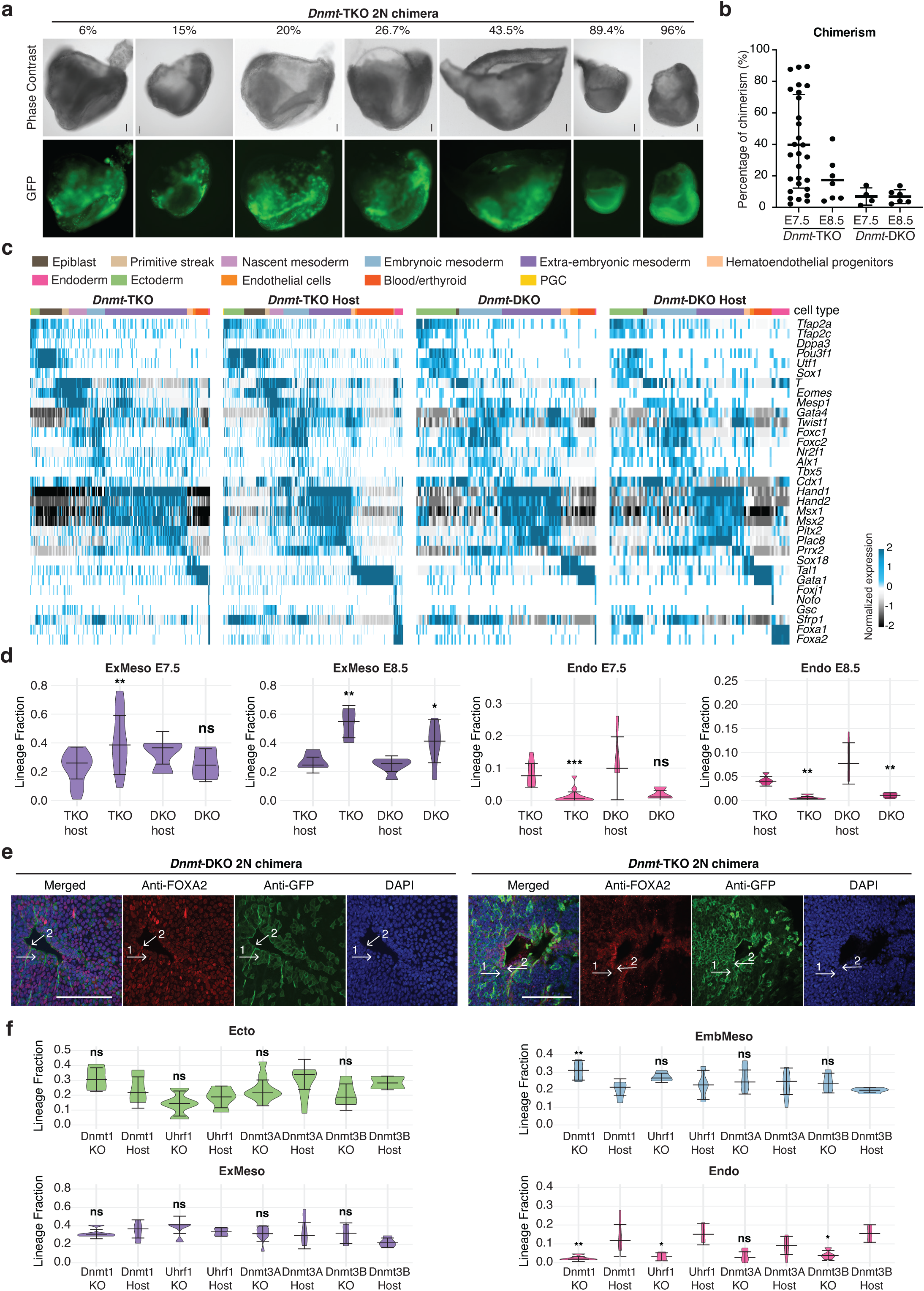
*Dnmt*-TKO and -DKO differentiation in the context of mixed chimeras. **a**, Images of *Dnmt*-TKO mixed chimeras with variable chimerism. *Dnmt*-TKO cell percentage per embryo analyzed by flow cell cytometry was marked on top. All embryos are oriented with the anterior part to the left. Scale bars, 100 μm. **b**, Flow cytometric analysis for the degree of chimerism per embryo, as measured by the % of GFP-positive cells. Data are represented as mean ± SD. Note that chimerism cannot be directly compared between different genotypes in this plot since very few DKO embryos were analyzed. **c**, Heatmap showing enrichment (expression relative to the median across cell states) of marker gene expression (rows) across metacells (columns) from indicated genotypes, colored by annotated cell states (top). **d**, Quantification of extraembryonic mesoderm (ExMeso) frequency in *Dnmt*-TKO and *Dnmt*-DKO mixed chimeric embryos isolated at E7.5 and E8.5. ns, not significant; *, *p* value < 0.05; **, *p* value < 0.01; ***, *p* value < 0.001. **e**, Images of early to late head-fold stage *Dnmt*-DKO and -TKO mixed chimeric embryo sections stained with anti-FOXA2 (red), anti-GFP (green, refers to mutant cells), and DAPI (blue). Arrow shows cells with co-localization of FOXA2-positive cells with *Dnmt* mutant. Scare bars, 100 μm. **f**, Quantification of the frequency of major lineages per indicated genotype. Shown are comparisons between host and mutant cell contributions. ns, not significant; *, p value < 0.05; **, *p* value < 0.01.

**Extended Data Fig.4.**
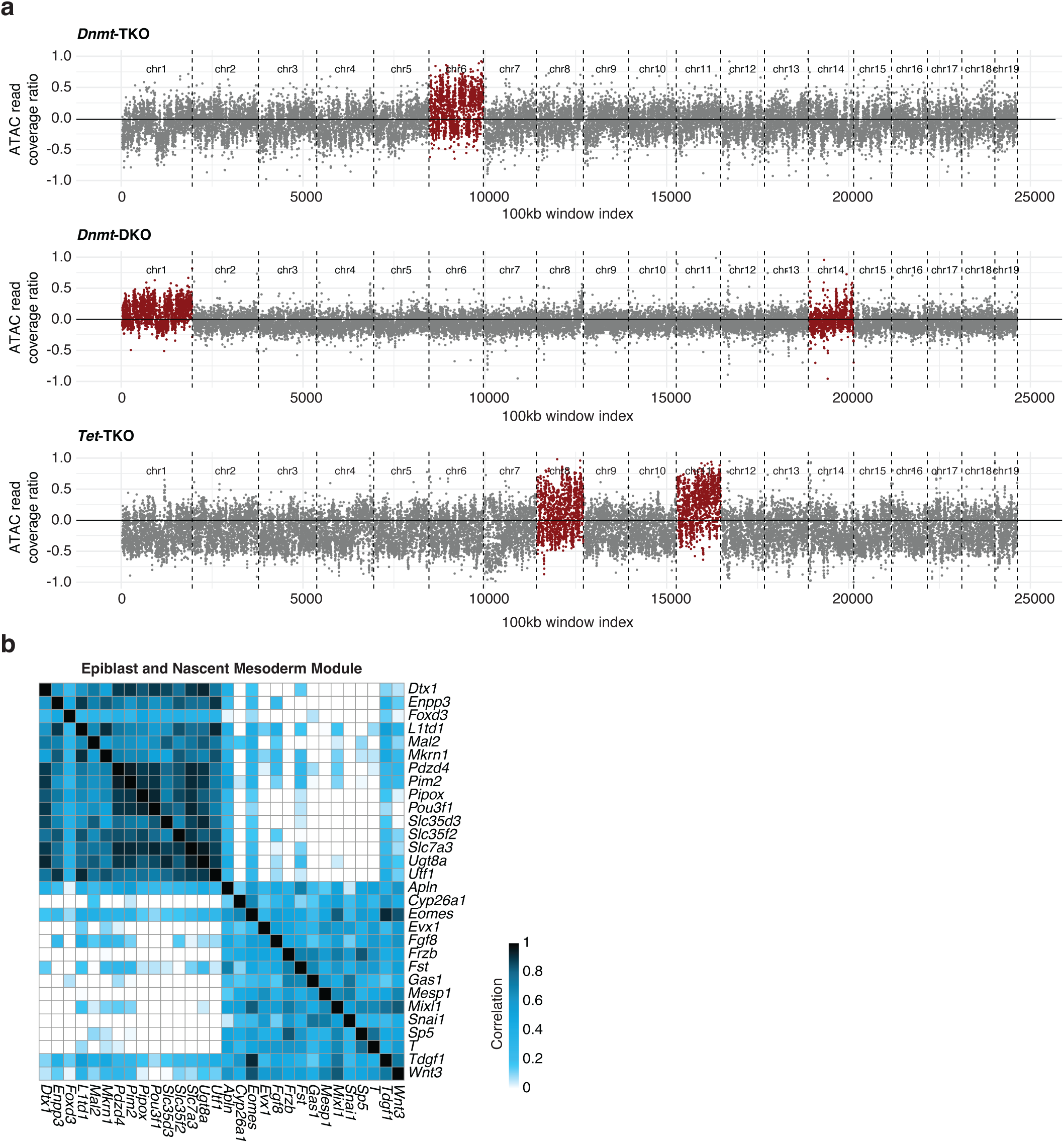
Gene exclusion based on chromosomal abnormalities and definition of cell state modules. **a**, Comparison of ATAC-seq coverage per 500 bp along the genome between mutants and host cells. For each bin, coverage is normalized by the mean coverage per bin in the corresponding genotype, and the y-axis shows the ratio of normalized mutant coverage to normalized host coverage. Chromosomes identified as trisomic are highlighted in red. **b**, Correlation heatmap depicting WT expression of epiblast and nascent mesoderm module genes. Expression for each gene was extracted from a previously established single-cell atlas of WT embryos^47^. See Methods for details.

**Extended Data Fig.5.**
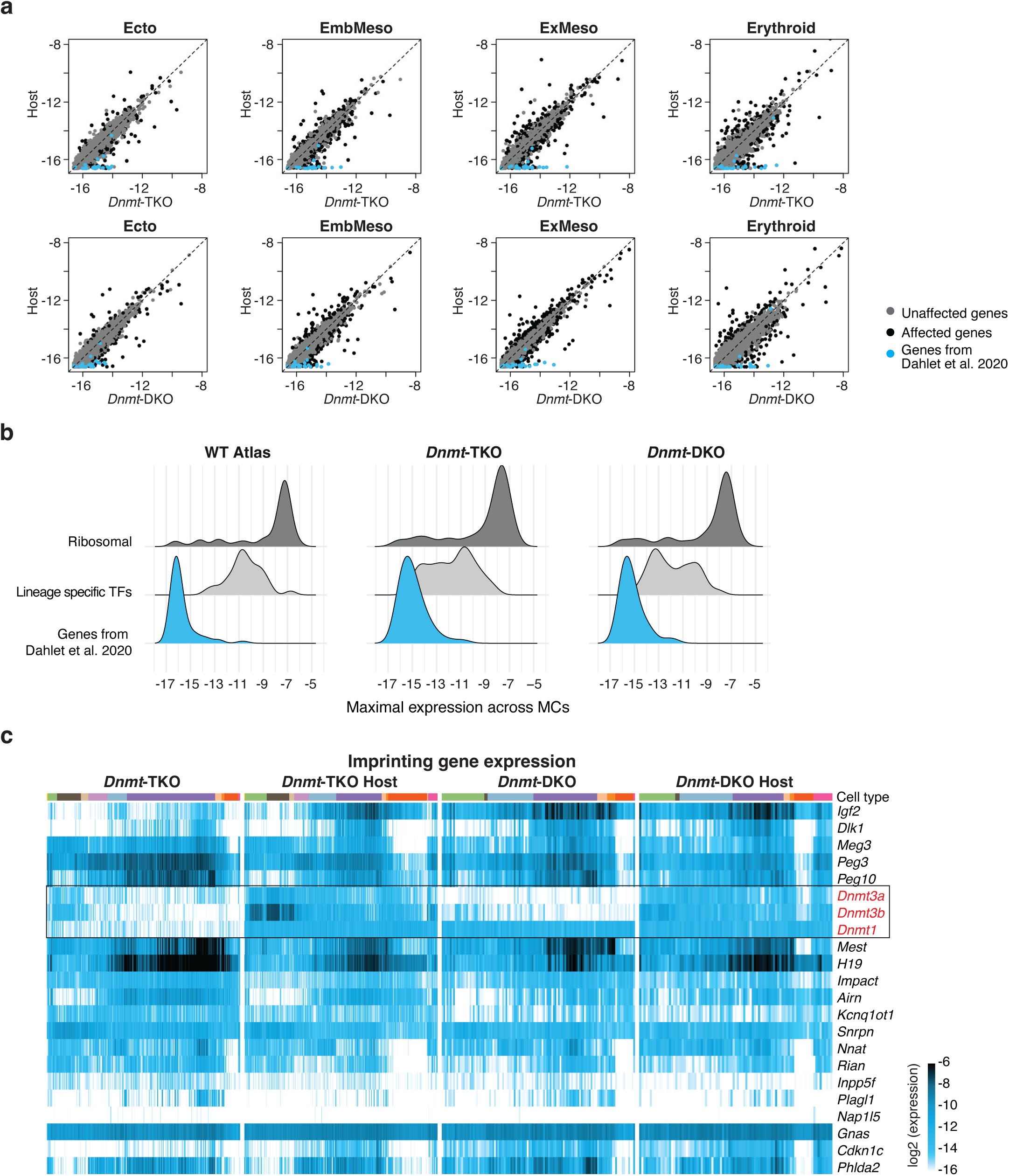
Global gene expression changes in mixed chimeric context after *Dnmt* depletion. **a**, Comparison of gene expression between *Dnmt*-TKO, -DKO and their respective host cells in different lineages. Previously reported methylation-affected genes from Dahalet et al., 2020 are highlighted in light blue; Significantly skewed genes (*q* val < 0.01) are highlighted in black (see Supplementary Table 1). **b**, Metacell maximal expression distributions for ribosomal genes, lineage-specific TF genes, and previously reported genes from Dahalet et al., 2020, in WT, *Dnmt*-TKO, and *Dnmt*-DKO embryos. **c**, Heatmaps depicting absolute expression (log2 of UMI frequency) of imprinted genes per metacell (columns) across mutants and their respective hosts, colored by annotated cell types (top). Knocked-out genes (*Dnmt*1, *Dnmt*3a, *Dnmt*3b) are highlighted in red for reference.

**Extended Data Fig.6.**
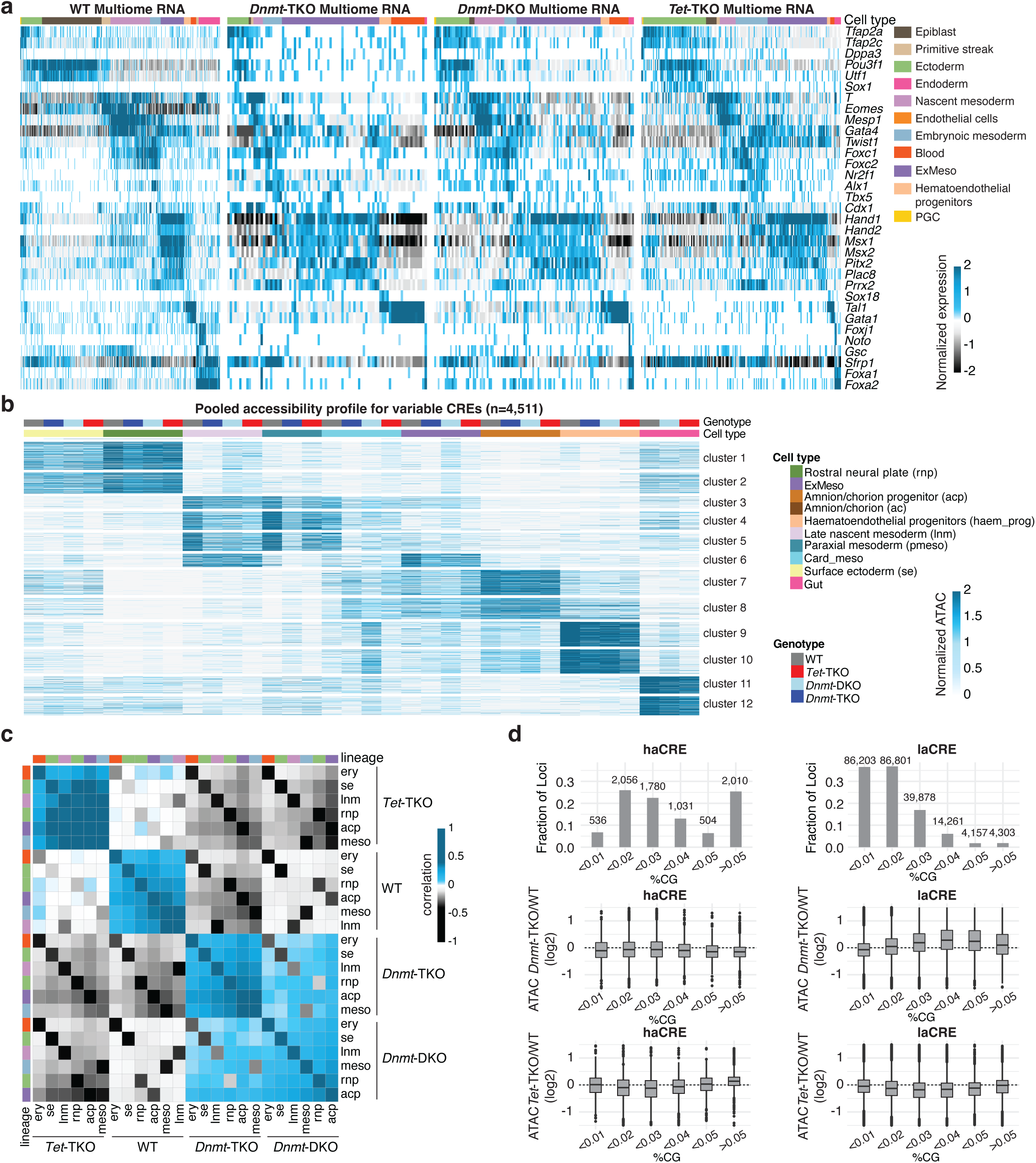
RNA and chromatin accessibility profiling of *Dnmt*-TKO and *Dnmt*-DKO cells. **a**, Heatmap showing enrichment of marker gene expression (rows) across metacells (columns) from indicated genotypes in the multiome assay, colored by annotated cell states (top). **b**, Clustered heatmap showing accessibility profiles across pooled cell states per lineage (see annotations on the right). Loci were chosen based on their preference for a specific cell state in the WT atlas (across rows, log2(maximum expression/median expression) > 2, logged maximum expression > -15). **c**, Auto-correlation matrix for mean-normalized ATAC profiles of different lineages and cell states. **d**, Top panel: Fraction of high- and low-accessibility CREs within each CpG content bin (see Fig. 3c). Absolute numbers of regions in each bin are indicated. Middle and bottom panels: Changes in chromatin accessibility of these elements in *Dnmt*-TKO (middle) and *Tet*-TKO (bottom) relative to WT. Data are shown for the extraembryonic mesoderm lineage.

**Extended Data Fig.7.**
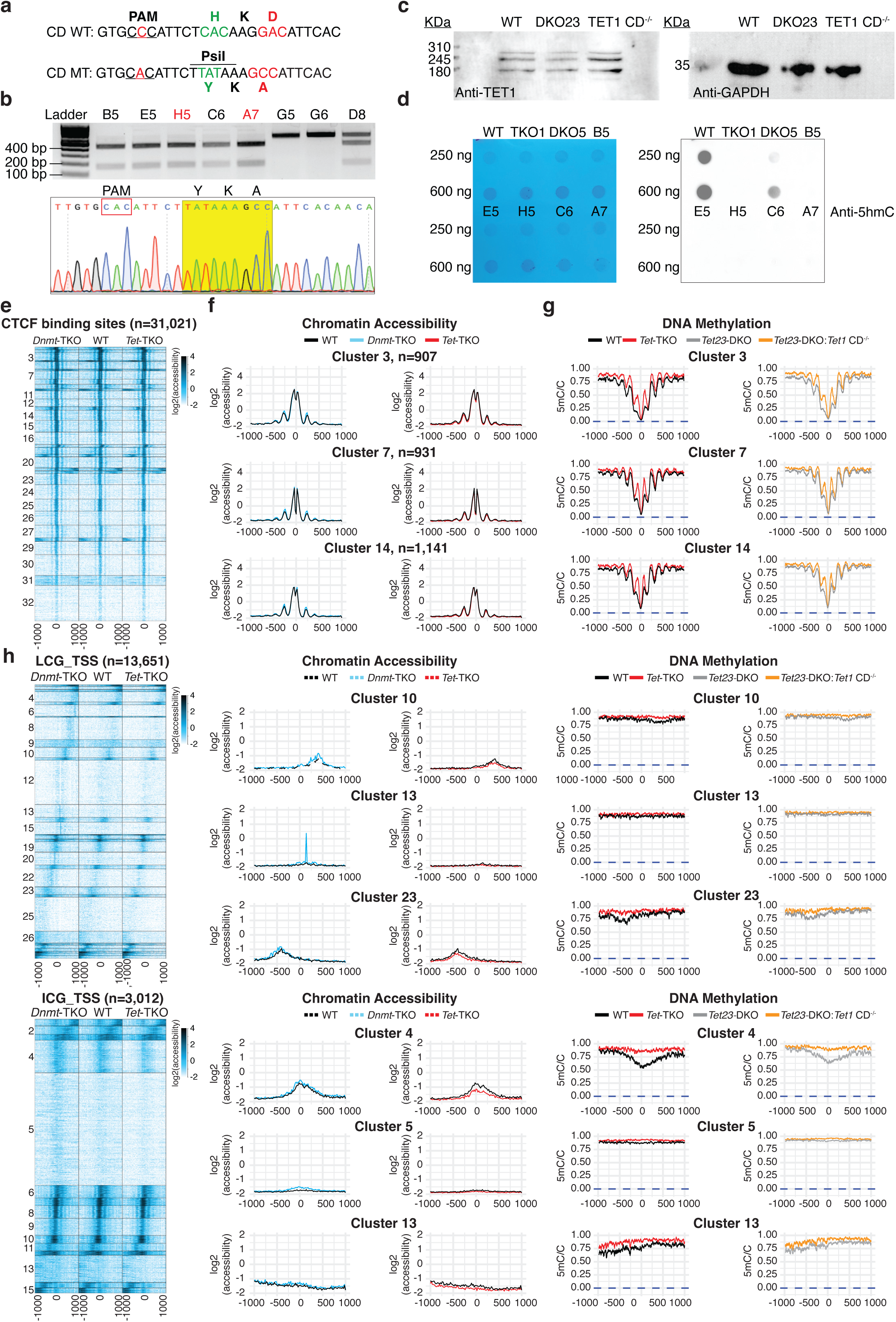
Impact of *Dnmt*-TKO and *Dnmt*-DKO on chromatin accessibility for CTCF binding sites, LCG_TSS and ICG_TSS. **a**, Mutation strategy for the generation of TET1 catalytic mutant. **b**, Gel electrophoresis and Sanger validation of candidate TET1 catalytic mutant. PCR products using primers flanking the mutation site were restricted with *PsiI* and assessed with gel electrophoresis. The untargeted WT allele is 614 bp, and the targeted allele was digested into a 439 bp and a 175 bp band, respectively (upper panel). Clones with two digested bands were further subjected to Sanger sequencing, and the result of a representative mutant clone is shown (lower panel). Note that the original PAM sequence was mutated (marked in yellow) to prevent repeated cutting by the Cas9 protein. Clones labeled in red were chosen for chimeric injections. **c**, Western blot validation for TET1 catalytic mutant in *Tet23*-DKO mutant background. **d**, Dot blot validation for TET1 catalytic mutant in *Tet23*-DKO mutant background. **e**, Heatmap of clustered chromatin accessibility profiles for 2kb loci (rows) centered and oriented according to CTCF binding sites, in *Dnmt*-TKO (left), WT (center), and *Tet*-TKO (right), ectoderm profiles. **f**, Quantitative zoom-in for representative clusters from **e**, comparing the three genotypes. **g**, As in **f**, but showing DNA methylation profiles in WT and three different TET mutant lines. The Y-axis represents averaged methylation, ranging from 1 (fully methylated) to 0 (unmethylated). Blue dashed lines indicate the zero expected methylation levels for *Dnmt*-TKO. **h**, Clusters of chromatin accessibility profiles for 2k-bp windows surrounding low CG TSSs (CpG density < 2%) and intermediate CpG TSSs (2% < CpG density < 4%). Chromatin accessibility and methylation metadata profiles are also shown for representative clusters (see main Fig.4).

**Extended Data Fig.8.**
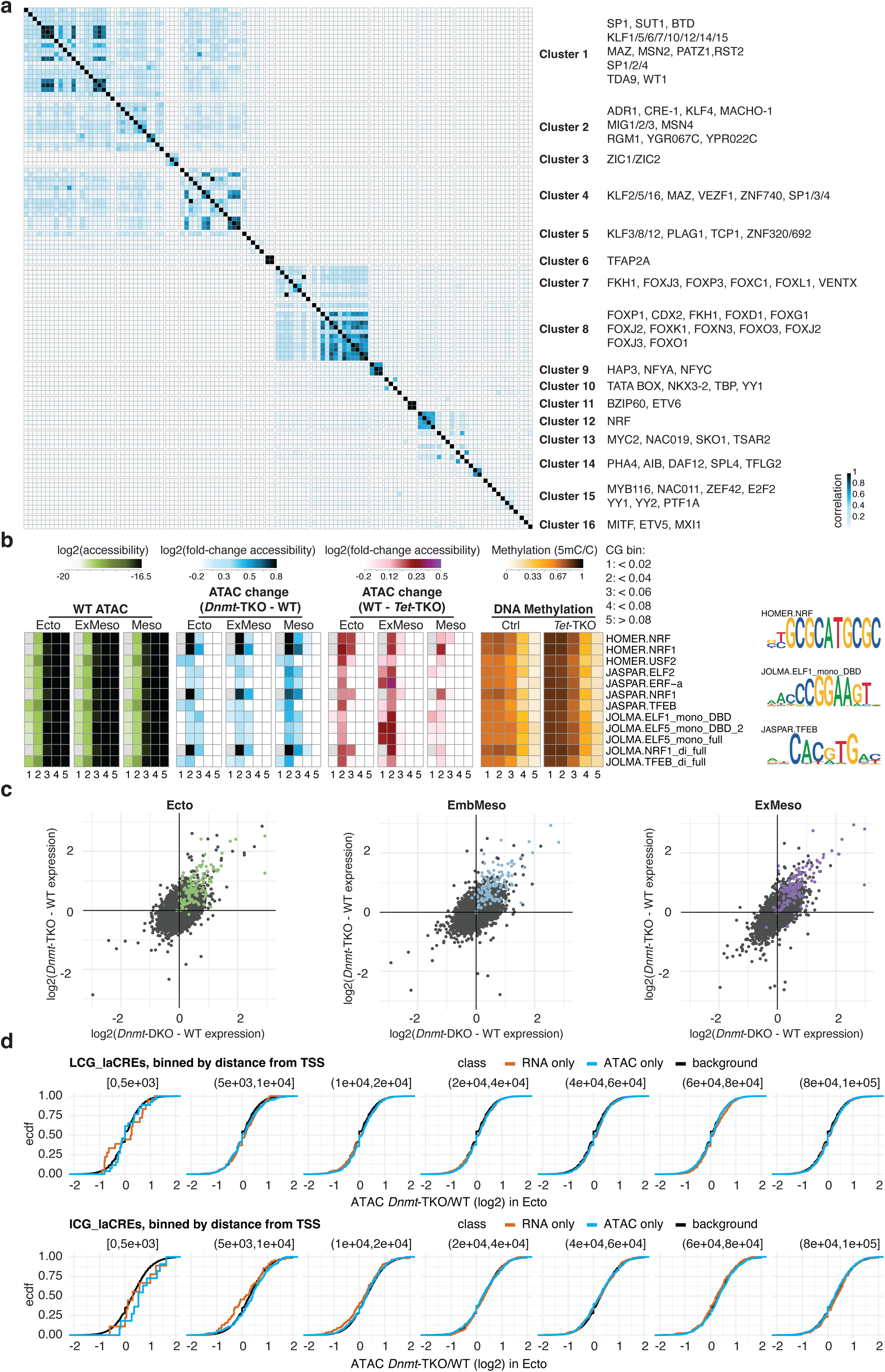
Impact of *Dnmt*-TKO and *Dnmt*-DKO on gene expression and putative TF binding at TSS. **a**, Co-occurrence similarity matrix (using Jaccard similarity) comparing the loci composition of motif-favoring clusters. The motif order per group corresponds to the row order in Fig.5a. **b**, A repetition of Fig.5a analysis for TSS loci only. CpG content bins are partitioned at [0-2%, 2%-4%, 4%-6%, 6%-8%, >8%]. Representative motif logos for each cluster are depicted to the right. **c**, Comparison of log2(*Dnmt*-TKO/host) vs. log2(*Dnmt*-DKO/host) gene expression ratios for all genes, with genes associated with both ATAC and RNA methylation-sensitive TSSs (see Fig.5c) highlighted in color. Analysis was repeated for three cell states: Ecto, EmbMeso, and ExMeso. **d**, Distributions of log fold change accessibility for *Dnmt*-TKO vs. WT calculated for neighboring CREs in respect to the TSSs highlighted in Fig.5c (belonging either to ICG_laCREs or LCG_laCREs category). CREs are binned by distance from a highlighted TSS. Background distribution is represented by log fold change accessibility for *Dnmt*-TKO vs. WT for all CREs of either the ICG_laCREs or LCG_laCREs category.

**Extended Data Fig.9.**
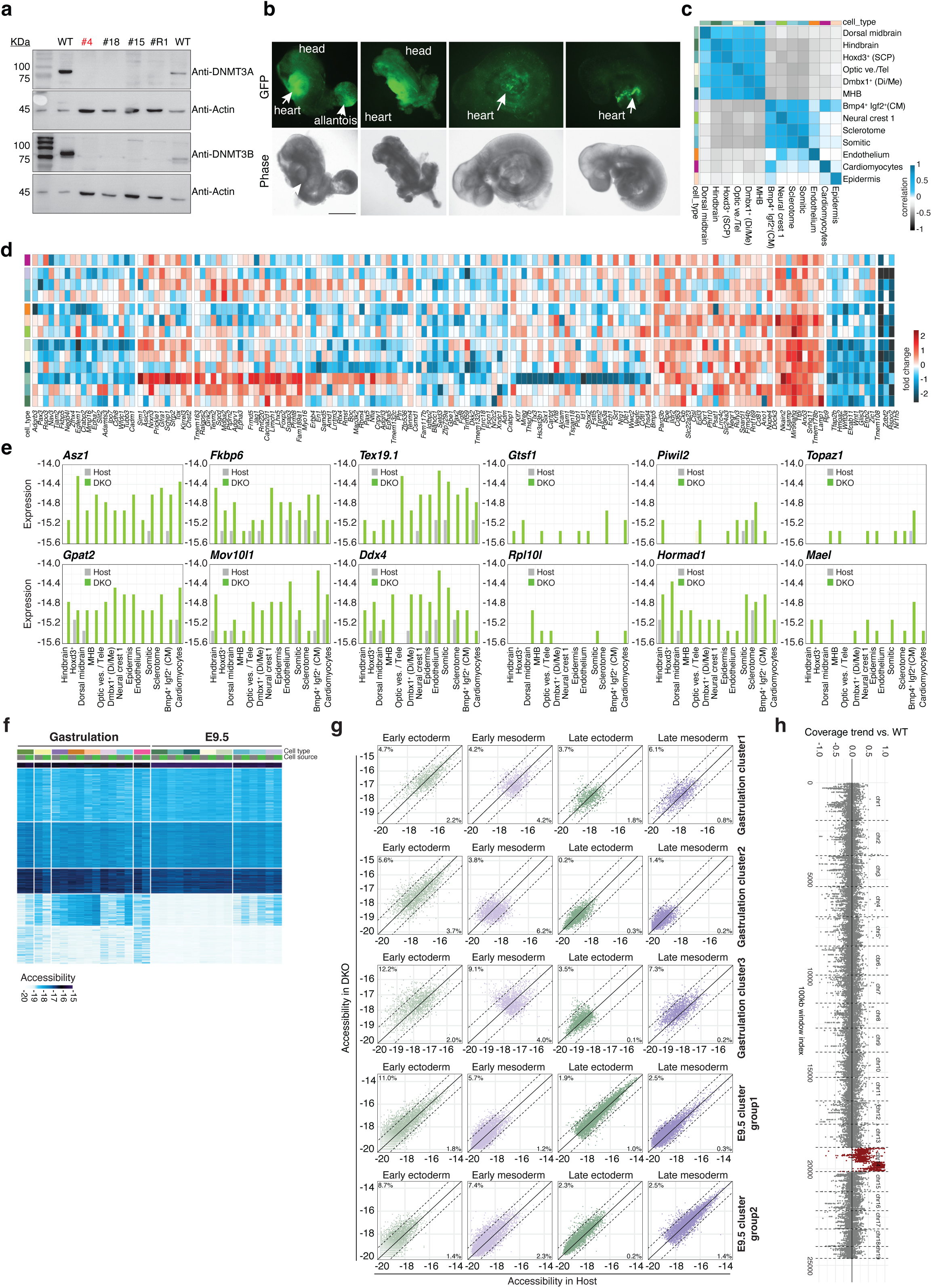
Effects of DNA methylation loss on transcription and chromatin accessibility during early organogenesis. **a**, Western blot validation for a new *Dnmt*-DKO mESCs clone. The clone labeled in red was used for subsequent chimeric injections. *Dnmt*-DKO clone #18, *Dnmt*-TKO clone #15, and R1 were used as controls. **b**, Representative images of relatively high and low contribution *Dnmt*-DKO chimeras at E9.5. Scale bar, 750 μm. **c**, Heatmap matrix showing pooled expression correlation between E9.5 cell states. Rows represent *Dnmt*-DKO expression, while columns represent host expression. **d**, Differential gene expression across properly sampled E9.5 cell states (total UMIs > 250,000 for both host and *Dnmt-*DKO genotypes). Values represent log fold-change (DKO vs. host) per cell state. **e**, Absolute expression of germline-associated genes across E9.5 cell states in *Dnmt*-DKO cells and host cells, revealing ectopic germline gene expression in *Dnmt*-DKO cells. **f**, Clustered heatmap showing chromatin accessibility profiles across pooled cell states per lineage for all CREs fulfilling log2(*Dnmt*-DKO/host) > 1.5 for at least one cell state. Subsets of these clusters are displayed in Fig.6e. **g**, Scatter plots comparing chromatin accessibility between *Dnmt*-DKO (y-axis) and host (x-axis) cells across four groups of cell states (see Fig.6f for exact composition): early ectoderm, early mesoderm, late ectoderm, and late mesoderm (columns). Rows correspond to CRE categories defined by lineage and developmental stage (see Fig.6e). Solid diagonal denotes equal accessibility, and dashed lines indicate a two-fold change threshold. Percentages denote the fraction of CREs exceeding the two-fold threshold in each direction. **h**, Comparison of ATAC-seq coverage per 500 bp along the genome between mutants and host cells. For each bin, coverage is normalized by the mean coverage per bin in the corresponding genotype, and the y-axis shows the ratio of normalized mutant coverage to normalized host coverage. Chromosomes identified as trisomic are highlighted in red.

## Methods

### Culture of mESCs

All mouse embryonic stem cells (mESCs) were cultured on irradiated mouse embryonic fibroblasts (MEFs) with standard medium: 500 ml DMEM (gibco, 41965-039), 20% fetal bovine serum (Hyclone, SH30071.03), 10 mg recombinant leukemia inhibitory factor (LIF, homemade), 0.1 mM beta-mercaptoethanol (gibco, 31350-010), penicillin/streptomycin (Biological Industries, 03-031-1B), 1 mM L-glutamine (Biological Industries, 03-020-1B), and 1% nonessential amino acids (Biological Industries, 01-340-1B).

For chimera assays, mESCs were cultured in standard medium for 2-3 days and replaced with 2i+LIF medium containing an additional 1 mM PD0325901 (Sigma, PZ0162) and 3 mM CHIR99021 (Sigma, SML1046) for another growth of 10-12 h before injection.

### Generation of *Dnmt* KO mESCs

We first generated a WT cell line, named Ctrl4, by electroplating *Kpn*I linearized Rosa26 mT/mG plasmid (Addgene, #17787) into V6.5 mESCs (Jaenisch lab, CVCL_C865) under the following conditions: voltage: 500, capacitance: 25, Resistance: ∞, and cuvette: 4 (BioRad, Gene Pulser Xcell). Puromycin (2 mg/ml stock) was added at a 1:2000 ratio 24 h post-electroporation. Clones that survived a 6-day puromycin selection were picked up and further validated by PCR for correct integration using primers listed in Supplementary Table 4. Subsequently, Ctr4 and a previously reported WT cell line Ctrl3^48^ were used to generate *Dnmt*-TKO (Ctrl4 background) and *Dnmt*-DKO (Ctrl3 background) mESCs using the CRISPR/Cas9-mediated genome editing method.

Guide RNA (gRNA) targeting each *Dnmt* gene was designed using CRISPR online designer (http://crispor.tefor.net/). Detailed gRNA sequences are listed in Supplementary Table 4. Each gRNA was cloned into a px330 plasmid under the U6 promoter (Addgene plasmid #98750). According to the provider’s instructions, the correctly cloned plasmid was transfected into pre-plated Ctrl3 or Ctrl4 mESCs using TransIT-X2 Transfection Reagent (Mirus Bio, MIR6003). Cells were sorted for BFP 48 h post-transfection. Multiple single clones were picked up for further validation. One, two, or three sgRNAs were transfected to screen for *Dnmt* single, double, and triple KO mESCs separately.

The validated *Dnmt*-TKO clones were further treated by Cre recombinase protein to convert the membrane tdTomato into membrane GFP for better visualization using an inverted fluorescent microscope.

### Generation of TET1 catalytic domain mutant ESCs

To test if the demethylase activity of TET1 protein is needed during early gastrulation, we used single-strand oligo donor (ssODN) mediated-CRISPR/Cas9 to introduce point mutations to the catalytic domain of TET1 protein in *Tet2* and *Tet3* DKO cell line DKO23#5^48^. Specifically, the ssODN harbors mutations of H1652Y and D1654A, which will also introduce the recognition site of restriction enzyme *PsiI* if correctly targeted. A silent mutation within the PAM sequence was introduced to prevent Cas9 from cleaving the targeted site after the desired modification is achieved (Supplementary Table 4). The final 97 bp ssODN was purchased as Alt-R HDR Donor oligo from IDT. Additionally, guide RNA with its PAM sequence upstream of the mutation sites was cloned into the px-330 BFP plasmid (Supplementary Table 4).

1 µl of 100 µM ssODN and 1 µg guide RNA-px330 plasmid were co-transfected into DKO23 cell line using TransIT-X2 Transfection Reagent (Mirus Bio, MIR6003). Primers flanking the mutation sites were used to amplify fragments from individual clones (Supplementary Table 4). Afterward, 5-6 µl PCR products were digested in a 30 µl solution with 0.5 µl *PsiI* enzyme (NEB, R0744) at 37°C overnight. When assessed by gel electrophoresis, correctly targeted clones will yield DNA bands of 175 bp and 439 bp. For clones with correct *PsiI* digestion, we purified the PCR product before digestion with homemade Ampure beads, and further validated the mutation by Sanger sequencing at the Weizmann Institute of Science.

### Chimera and Tetraploid Complementary Assay

All animal procedures were approved by the Institutional Animal Care and Use Committee and were performed in strict adherence to Weizmann Institute guidelines. Mice were monitored for health and activity and were given ad libitum access to water and standard mouse chow with 12 h light/dark cycles.

To generate chimera embryos, KO mESCs were injected into diploid or tetraploid B6D2F x B6D2F1 E3.5 blastocysts and surgically implanted into E2.5 postcoital pseudo-pregnant Hsd:ICR (CD-1) females following standard procedures. Embryos were harvested and dissected in ice-cold 1×PBS, followed by imaging in a drop of DMEM medium supplemented with 10% fetal bovine serum under a Nikon Eclipse Ti2 inverted microscope. Embryos positive for KO cells (GFP positive) were trypsinized at 37°C for 5 min and quenched with 4 times the volume of sorting buffer (1×PBS with 0.5% BSA) and kept on ice until flow cell cytometry (BD FACSAria™ and FACSymphony S6). Mixed chimeric embryos with either too high or too low (sparse) contribution of injected cells (based on the fluorescent signal) were excluded from index-sorting, given that they wouldn’t provide enough control/host or KO cells for single-embryo resolution.

### Dot blot for 5mC and 5hmC level detection

To screen for potential KO clones after CRISPR/Cas9-mediated gene editing, genomic DNA was extracted from every single clone after 5-6 passages post-transfection. A total of 50-100 ng genomic DNA was blotted onto nitrocellulose membranes with the Bio-Dot microfiltration apparatus (Bio-Rad, 1706545). Afterward, the membrane was air-dried for 20 minutes and cross-linked on each side for 5 minutes with a UV-agarose gel box. 5% non-fat milk in 1x PBST was used to block the membrane, followed by primary and secondary antibody incubation. Antibodies used for this assay: anti-5mC (Abcam, ab10805), anti-5hmC (Epigentek, A-1018), Goat anti-Mouse IgG (H+L), HRP (Invitrogen, 31430). Signal was developed with SuperSignal™ West Pico PLUS Chemiluminescent Substrate (Thermo, 34579), and images were taken using ChemiDoc^TM^ MP Imaging System (BioRad).

### Western blot for protein level detection

For the detection of DNMT1, DNMT3A2, DNMT3B, DNMT1, UHRF1, and TET1 proteins, mESCs cultured in standard mESCs medium were harvested after MEF depletion. For the detection of DNMT3A1, 9-day EBs were collected after differentiation in MEF medium (supplemented with 10% serum, without 2i and LiF) in a petri dish. All the pellets were lysed in lysis buffer (150 mM sodium chloride, 1% Triton x-100, 50 mM Tris HCl pH8.0, with freshly added proteinase inhibitor) for 30 min on ice. The supernatant was reserved for total protein quantification using the Pierce BCA protein assay kit (Thermo Fisher Scientific, 23227). A total of 10-20 µg proteins were loaded for SDS-PAGE analysis. To re-probe the same blotted membrane with other primary antibodies, the membrane was immersed in stripping buffer (2% SDS, 0.0625 M Tris HCl, pH 6.8, 0.008% ß-mercaptoethanol) at 50°C for 45 min with gentle shaking. After intensive wash with 1xPBS, the membrane was blocked in blocking buffer (5% non-fat milk in PBST), followed by primary antibody probing. Primary antibodies used in the assay: anti-DNMT3A (NB120-13888, Novus Biologicals), anti-DNMT3B (NB300-516, Novus Biologicals), anti-DNMT1 (Cell Signaling, 5032), anti-UHRF1 (Santa Cruz, sc-373750), anti-TET1 (GeneTex, GTX124207), anti-alpha Tubulin (Millipore, ABT170), anti-GAPDH (Abcam, ab181602, Proteintech, 60004-1-lg). Secondary antibodies used: Goat anti-Rabbit IgG (H+L), HRP (Invitrogen, 31460), and Goat anti-Mouse IgG (H+L), HRP (Invitrogen, 31430). Signal was developed with SuperSignal™ West Pico PLUS Chemiluminescent Substrate (Thermo, 34580), and images were taken using ChemiDocTM MP Imaging System (Bio-Rad).

### Whole-mount immunofluorescence and confocal imaging

To detect the contribution of *Dnmt*-DKO and -TKO mESCs into embryonic endoderm lineage, we harvested *Dnmt*-DKO and -TKO 2N chimeric embryos at E7.5 and fixed them in 4% paraformaldehyde at 4°C overnight. The staining procedure was performed as described^48^. Primary antibodies used: anti-FOXA2 (Abcam, ab108422), anti-GFP (Abcam, ab13970), anti-mCherry (SICGEN, AB0040-200).

After the staining procedure, embryos were unfolded with forceps from the lateral side, placed onto a pre-treated glass slide with their ventral side facing up, mounted with a drop of mounting medium, then sealed with a coverslip^69^. Z-stack images were acquired with a Zeiss LSM 900 inverted confocal microscope (Carl Zeiss). Co-localization of FOXA2-positive cells and mutant cells (GFP-positive) was analyzed in ImageJ software, and images presented are max projection plots.

### Nuclei isolation for multiome sequencing

Nuclei for the multiome assay were prepared from Hsd:ICR (CD-1) embryos or cells sorted from chimeric embryos based on the experiment type. For the E9.5 multiome experiment, instead of taking the whole mixed chimeric embryo for single cell suspension preparation, we dissected out the head and heart regions from the chimera embryos and sorted mutant and host cells for the downstream multiome assay. Single nuclei suspension was prepared following the demonstrated “low cell input nuclei isolation” protocol from 10x Genomics with minor changes. Briefly, cells were washed once with 500 µl cold PBS/0.04% BSA and resuspended in 50 µl PBS/0.04% BSA in a 200 µl DNase/RNase-free tube. Cells were then pelleted at 300 rcf for 5 min at 4°C. After removing the supernatant, cells were lysed in 45 µl chilled lysis buffer (10 mM Tris-HCl, pH7.4, 10 mM NaCl, 3 mM MgCl_2_, 0.1% Tween-20, 0.1% Nonident P40 substitute, 0.01% digitonin, 1% BSA, 1 mM DTT) with freshly added RNase inhibitor (final concentration 1U/µl, Sigma, 3335399001) for 3 min on ice. Afterward, 100 µl chilled wash buffer (10 mM Tris-HCl, pH 7.4, 10 mM NaCl, 3 mM MgCl_2_, 1% BSA, 0.1% Tween-20, 1mM DTT, 1U/µl RNase inhibitor) was added directly to the tube, and mixed by another 5 times gentle pipetting. The nuclei were pelleted at 500 rcf for 5 min at 4°C. After removing 145 µl supernatant, 45 µl chilled diluted nuclei buffer (10x Genomics) was added to the tube without dislodging the pellet. Followed by a centrifuge at 500 rcf for 5 min at 4°C, the final nuclei were resuspended in 7 µl diluted nuclei buffer. 2 µl nuclei suspension was taken to quantify the concentration with a Countess II (Thermo) and to check the quality under an inverted microscope. >95% of nuclei stained positive for trypan blue, and the nuclei were found to have the expected morphology. Nuclei were further diluted to the suggested concentration range suitable for preparing a 4000-6000 nuclei library if needed.

### Multiome library preparation

Single nuclei suspensions were loaded on Chromium Next GEM Chip J (10x Genomics) to generate single-cell GEMs. Single-cell ATAC library and gene expression library were constructed separately following the manufacturer’s instructions. Library size distribution and abundance were assessed with a D1000 or D5000 ScreenTape (Agilent), and their concentration was quantified with a Qubit 4 Fluorometer (Thermo). Libraries were sequenced on an Illumina NovaSeq 6000 instrument in paired-end mode. Sequencing settings are: ATAC library: read1: 50 cycles, index1: 8 cycles, index2: 24 cycles, read2: 49 cycles. RNA library: read1:28 cycles, index1:10 cycles, index2: 10 cycles, read2: 90 cycles. E9.5 multiome libraries are sequenced on an Illumina NovaSeq X Plus instrument in paired-end mode. Settings are: ATAC library: read1: 151 cycles, index1: 10 cycles, index2: 24 cycles, read2: 151 cycles. RNA library: read1:151 cycles, index1:10 cycles, index2: 10 cycles, read2: 151 cycles.

### Enzymatic Methyl-seq (EM-seq)

*Tet1*CD^-/-^, *Tet*2 and *Tet3* DKO23, and Ctrl3 mESC were separately injected into diploid B6D2F1 E3.5 blastocysts and allowed to develop in pseudo-pregnant ICR females. Afterward, cells with indicated genotypes were sorted using flow cytometry from streak to late bud and headfold stage chimeric embryos. Genomic DNA was made from those cells using Quick-DNA^TM^ miniprep plus kit (Zymo Research, D4068). For EM-seq library preparation, 10-50 ng genomic DNA was taken from each sample and mixed with methylated lambda DNA and unmethylated pUC19 DNA at a 1:100 ratio in a 0.5 ml tube. After cooling on ice for 10 min, DNA was sheared using the Diagenode Bioruptor 300 with the following settings: Energy: High, 30 s ON, and 30 s OFF for 10 cycles. Sonication was repeated 3 times (30 cycles in total). Subsequently, 1µl sonicated DNA was assessed using TapeStation 4150 (Agilent). A sample with a DNA size of around 250 bp proceeded to library preparation using NEBNext® enzymatic methyl-seq kit (New England Biolabs, E7120S) according to the manufacturer’s instructions. In brief, sheared DNA was end-repaired and ligated with EM-seq adaptors. After purification, DNA was oxidized by TET2 protein, followed by DNA denaturing using formamide (Ambion, AM9342). Subsequently, purified DNA was deaminated by APOBEC protein. The deaminated DNA was then purified and amplified with a specific EM-seq index primer. Tagged DNA products were cleaned and further assessed by Qubit 4 Fluorometer and TapeStation 4150. Finally, libraries were pooled and sequenced on Illumina NovaSeq X Plus with the following parameters: read1:150 cycles, read2: 150 cycles, index1: 17 cycles, index2: 8 cycles.

### Single-cell transcriptional library preparation

Single-cell cDNA libraries were prepared using the massively parallel single-cell RNA-seq (MARS-Seq) method as described^70^. Briefly, single cells were sorted into 384-well cell capture plates containing 2 µl of lysis solution and barcoded poly (T) reverse-transcription (RT) primers. 0.5 µl RT buffer was added to each well by MANTIS microfluidic liquid handler (FORMULATRIX) to get complementary DNA (cDNA), followed by 1 µl Exonuclease I mixture treatment without heat inactivation. Afterward, all liquid from each 384-well plate was pooled into the VBLOCK200 reservoir (Clickbio) by centrifuge, thus denoted as one library. The library was converted to double-stranded DNA, transcribed using T7 RNA polymerase, and fragmented. Following this, a unique plate adaptor was added to each library to be sequenced together, and RNA::ssDNA (plate adaptor) ligation was performed. The purified ligates were applied to a second RT step with a primer complementary to the ligated adaptor using Affinity Scripts RT enzyme, and the library was completed by cDNA enrichment through PCR amplification. The library concentration and profile were evaluated before sequencing.

### Metacell analysis of MARS-seq data

First, we filtered out cells outside a umi count range of 1,000 to 20,000. In addition, we used our WT atlas to identify genes associated with high expression in the Visceral endoderm, Anterior visceral endoderm, Parietal endoderm, and Extraembryonic ectoderm, forming gene sets for each such type. We filtered out cells that had high summed expression (see code companion for thresholds) of these modules over the overall expression distribution. Then, we ran the Metacells algorithm^52^ on filtered count matrices, separately for knockouts and hosts in each mutant type, to generate individual metacell models. In all metacell runs, we employed the same parameters (target size = 50 cells per metacell) and gene sets (excluded, lateral). To annotate metacell models, we utilized the MCProj package^71^, which assigns metacells “projected states” based on transcriptional similarity with annotated metacells in a WT atlas. To simplify and enhance statistical analysis in downstream modeling, we formed coarse-grained types from cell states/types as specified in Supplementary Table 5.

### Cell state integrity analysis using gene programs in Fig.2

To assess cell state transcriptional integrity, we used the epiblast and nascent mesoderm gene sets defined in our previous work^48^. The epiblast module consists of genes that are highly correlated with *Utf1* expression: *Dtx1, Enpp3, Foxd3, L1td1, Mal2, Mkrn1, Pdzd4, Pim2, Pipox, Pou3f1, Slc35d3, Slc35f2, Slc7a3, Ugt8a*, and *Utf1*. The nascent mesoderm list consists of genes most highly expressed in the nascent mesoderm: *Apln, Cyp26a1, Eomes, Evx1, Fgf8, Frzb, Fst, Gas1, Mesp1, Mixl1, Snai1, Sp5, T, Tdgf1*, and *Wnt3*. Gene module scores were computed as the total UMIs for the gene set in single cells divided by the cells’ total UMI count.

### Divergence score assignment to individual embryos

We defined for each embryo and cell state of interest a gene expression profile computed using pooling RNA counts over all annotated cells from the respective embryo. We paired each query embryo to the atlas embryo with maximal correlation (*r_max_*). Divergence scores for a set of mutants were then defined as 1- *r_max_*. For control, we used a similar analysis applied to the paired atlas embryos but matched these to the reference atlas from which these control embryos were removed. Correlations were computed across a gene set of 887 genes showing high variance-to-mean ratio^53^, excluding genes located in trisomic chromosomes, X-linked genes, imprinted genes, and DNMT genes.

### Atlas-based differential mutant expression analysis in Fig.2f, g

We used MCProj also to derive the expected gene expression profile for each query metacell. Differential query/atlas expression was then defined by comparing pooled query expression (e.g. in a cell state) and the respective projected expression.

### Metacell analysis of RNA from 10X multiome data

RNA Count matrices generated by the 10X pipeline were processed using Metacells and MCproj as described above for MARS-seq. To account for systematic biases between MARS-seq and multiome 10X v3 chemistry, we applied the MCproj *project_correction* parameter to derive atlas annotations.

### Analysis of ATAC profiles and peaks

10X-mapped ATAC cells were grouped according to cell annotations derived through RNA analysis. Reads from groups of cells were then combined to generate cell state-specific accessibility tracks at 20bp resolution through the Misha package (https://tanaylab.github.io/misha/). A track combining all reads was also generated. We repeated this procedure for WT and mutant (*Dnmt*-TKO, *Dnmt*-DKO, and *Tet*-TKO) lines.

Our basic ATAC signal was considered as the total coverage in running 300bp windows. We computed a regional normalization track from the (300bp running window) total coverage track by computing truncated mean coverage in a larger running window of 20kb with a threshold at the 95^th^ coverage percentile. We then screened for raw peaks, which were defined as continuous intervals in which the ratio between 300bp coverage and regional 20k normalized coverage was at the top 2 genome-wide percentiles. After repeating this strategy for each dataset (WT and mutants), the final set of peaks for analysis was obtained by merging (unifying overlapping intervals) the four peak sets. This resulted in a set of 270,005 elements for downstream analysis.

### Correcting coverage and copy number effects in mutants’ ATAC analysis

We used the total WT ATAC coverage distribution over chromosomal bins to compute a reference coverage model. Copy number aberrations (CNAs) in mutants (in particular trisomies) were then defined as regions in chromosomes for which the median ratio between normalized mutant coverage and the reference coverage model showed a consistent skew. In practice, as highlighted in Extended Data Fig.4a and 9h, we detected trisomies in chr1 and/or chr14 for *Dnmt*-DKO, a trisomy in chr6 for *Dnmt*-TKO, and trisomies in chr8 and chr11 for *Tet*-TKO. More systematically, we computed for each chromosome *c* and mutant *line* a coverage ratio *cov_c,line_* as (*n_c,line_/n_line_): (n_c,wt_/ n_wt_*) (where n defines total read count in the respective chromosome and line).

We corrected for CNAs in all ATAC analyses by downsampling count matrices. Given any count statistics on peaks or regions in chromosome c and *line*, we transformed the data by a downsampling ratio of 1/*cov_c,line_*. This ensured balanced chromosomal mean coverage as well as balanced chromosomal accessibility variance in WT and mutants. For RNA analysis, we excluded genes on trisomic chromosomes when running differential expression analysis (Fig.2a-e, 5c).

### Normalizing ATAC accessibility intensities to a housekeeping standard

When comparing accessibility between lines and cell types, we further normalized ATAC count matrices to correct for the overall accessibility level of analyzed cells and the total sequencing depth of the dataset. For each line and coarse-grained cell type we computed the total ATAC coverage on TSSs of housekeeping genes (selected for simplicity as all genes coding for ribosomal proteins) 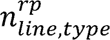. Given any count statistics on peaks or regions for a set of conditions 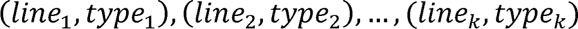, we identified the std 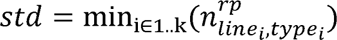 and downsampled all data by the ratios 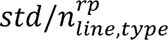. This ensured a similar coverage level for all conditions compared, thereby normalizing both mean and variance. We preferred this approach over normalization based on total coverage per se, as total coverage is skewed by additional technical and biological factors, while housekeeping genes accessibility can be assumed to be robustly constitutive. We note that this standardization is applied after the CNA corrections described above (where CNA normalization is specific to a line, and standardization is specific to each type, line combination).

### Analysis around CTCF binding sites in Extended Data Fig.7e-g

CTCF peaks were defined using published ChIP-seq data^72^ (identifying loci with coverage > 32). Peaks were extended by 100bp on each side, and the position showing maximum CTCF motif match in either the forward or reverse strand was identified and used to re-center peaks at the motif locus. We used a conservative definition of CTCF peaks for which almost all peaks contained a strong binding site. Weaker CTCF sites were analyzed as either TSSs or CREs.

### Definitions and classification of TSSs in Fig.3 and Fig.4

TSS coordinates were obtained from knownGene by UCSC Genome Database (https://hgdownload.cse.ucsc.edu/goldenpath/mm10/database/). ATAC peaks overlapping a TSS were merged into the TSS.

### Stratification of peaks by CpG content

We defined the CpG content of a genomic coordinate as the number of CpG dinucleotides in a 500bp window centered around the coordinate divided by 500. The CpG content of a peak is then defined as the maximal CpG content found within the peak. For controlling the CpG content effect, we either stratify by CpG content directly, or use an overall classification by CpG content over the ranges 0-2% (LCG), 2%-4% (ICG), >4% HCG.

### CREs accessibility intensity and using it for stratification of mutant effect in Fig.3g

In each line and cell type, following CNA normalization and coverage standardization, we compute the accessibility level of a peak relative to the total peak set as:

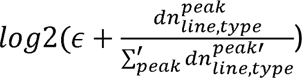

where *dn* is the standardized (downsampled) count matrix and ε= 10^-6^. ATAC intensities represented this way were typically distributed between -20 and -14 (see Fig.3f), and a threshold of -16.5 was used to classify high accessibility (ha) CREs from low accessibility (la) ones in downstream analyses that required grouping. We note that when grouping haCRE and laCREs (Fig.3c-e and Extended Data Fig.6d), we used the average intensity between WT ectoderm and extraembryonic mesoderm.

### Spatial clustering of loci within epigenetic hotspot categories

The spatial perspective for each hotspot was achieved by either centering its genomic interval around a local maximum in the CTCF signal (for CTCF classes), or a predefined TSS (for TSS classes), or by using the original center for the interval (for CRE classes). Once the interval center was defined, a genomic window of ±1kbp (2kbp total) was extended around it. These windows were further divided into 101 consecutive bins of equal size to capture chromatin accessibility patterns at high resolution. The resulting binned accessibility data for all hotspots were compiled into a matrix that was normalized (for CNA) and standardized (for coverage). We repeated matrix extraction and normalization for WT, *Dnmt*-TKO, and *Tet*-TKO and concatenated the trends for each locus. The combined matrix was then subjected to k-means clustering of loci using TGLKmeans (https://tanaylab.github.io/tglkmeans/). For visualization, we smoothed matrix columns using a window of 20 loci.

### EM-seq data analysis

EM-seq data were processed according to the NEBNext pipeline^73^. After observing that the samples from two time points (streak to late bud, headfold) produced indistinguishable methylation profiles, we averaged them together for each genotype and generated methylation tracks for all genotypes. We accumulated methylated and unmethylated reads to estimate average methylation trends in spatial clusters across different genotypes.

### Analysis of CRE motifs along with mutant accessibility effects

We used a position weight matrix library combining data from HOMER^74^, Jolma^75^, and JASPAR^76^. For each motif, we computed an approximated binding energy using the PREGO package (https://tanaylab.github.io/prego/) that summed up the high and low-affinity contributions of motifs for each sequence element. Binding energies estimation was performed on peak genomic intervals unified for 300bp in length.

Two types of ATAC-seq accessibility changes were analyzed: **log2(*Dnmt*-TKO) – log2(WT)** and **log2(WT) – log2(*Tet*-TKO)**. For each motif, binding energies were stratified into bins based on the CG content of the loci (<1%, <2%, <3%, <4%, <5%, >5%). Within each CG content bin, we compared accessibility changes between loci in the top 2% of motif binding energies and loci in the bottom 95%.

Motifs were classified as “motifs of interest” if they exhibited a Kolmogorov-Smirnov (KS) D statistic > 0.25 (FDR q < 10^-4^) for either the “up” (alternative = “less”) or “down” (alternative = “greater”) direction. The heatmap in Fig.5a highlights all motifs of interest.

### ATAC analysis over repeat families

The standard CellRanger pipeline excludes multi-mapped reads during processing, effectively removing most repeat elements from the final outputs. As a result, these outputs were unsuitable for repeat element analysis. To address this limitation, we utilized an intermediary BAM file output by CellRanger, specifically the “atac_possorted_bam.bam,” which still includes multi-mapped reads prior to filtering. Using this intermediary file, we constructed aggregated (marginal) coverage tracks for each experiment, capturing chromatin accessibility data prior to the exclusion of multi-mapped reads. These tracks allowed us to analyze the total coverage of repeat elements across different categories and experiments. Repeat elements were categorized and annotated based on UCSC RepeatMasker.

### Definition of methylation-sensitivity of genes in terms of ATAC and RNA

First, based on their proximity, we associated the ATAC signal at TSS loci with TSS coordinates annotated in the UCSC database. For cases where multiple loci matched the same gene name, we selected the locus with the highest mean ATAC signal in WT across three cell states: Ectoderm, Extraembryonic Mesoderm, and Mesoderm. A gene was deemed ATAC methylation-sensitive if its ATAC signal change, calculated as log_₂_(*Dnmt*-TKO) – log_₂_(WT), exceeded 0.8 in at least one of the three cell states.

To assess RNA methylation sensitivity, we utilized MARS-seq RNA expression data aggregated by cell state for the same three cell states. Genes were considered RNA methylation-sensitive if their RNA expression change, now calculated as log_₂_(mean *Dnmt*-KOs) – log_₂_(mean hosts), exceeded 1 in at least one cell state.

For both ATAC and RNA, we require an FDR of *q* < 0.01, considering testing all sufficiently covered (total > 20 downsampled UMIs for RNA, total > 10 downsampled UMIs for ATAC) TSSs/genes in three tests (one per cell state).

## Data availability

All datasets will be deposited in the Gene Expression Omnibus (GEO).

## Code availability

All code will be available on GitHub.

